# CD44 regulates epigenetic plasticity by mediating iron endocytosis

**DOI:** 10.1101/693424

**Authors:** Sebastian Müller, Fabien Sindikubwabo, Tatiana Cañeque, Anne Lafon, Antoine Versini, Bérangère Lombard, Damarys Loew, Adeline Durand, Céline Vallot, Sylvain Baulande, Nicolas Servant, Raphaël Rodriguez

## Abstract

CD44 is a transmembrane glycoprotein that is linked to various biological processes reliant on the epigenetic plasticity of cells, including development, inflammation, immune responses, wound healing and cancer progression. While thoroughly studied, functional regulatory roles of this so-called ‘cell surface marker’ remain elusive. Here, we report the discovery that CD44 mediates endocytosis of iron interacting with hyaluronates in tumorigenic cell lines and primary cancer cells. We found that this glycan-mediated iron endocytosis mechanism is enhanced during epithelial-mesenchymal transition, unlike the canonical transferrin-dependent pathway. This transition is further characterized by molecular changes required for iron-catalyzed oxidative demethylation of the repressive histone mark H3K9me2 that governs the expression of mesenchymal genes. *CD44* itself is transcriptionally regulated by nuclear iron, demonstrating a positive feedback loop, which is in contrast to the negative regulation of transferrin receptor by excess iron. Finally, we show that epigenetic plasticity can be altered by interfering with iron homeostasis using small molecules. This comprehensive study reveals an alternative iron uptake mechanism that prevails in the mesenchymal state of mammalian cells, illuminating a central role of iron as a rate-limiting regulator of epigenetic plasticity.

## INTRODUCTION

CD44 is a transmembrane glycoprotein that is implicated in many physiological and pathological processes including development and cancer progression among others (Ponta et al., 2003; Zöller, 2011). Some of these complex processes involve cells able to dynamically and reversibly shift between cell states through epithelial-mesenchymal transitions (EMT) (Brabletz et al., 2018; Nieto et al., 2016; Pastushenko et al., 2018). For instance, epithelial-mesenchymal plasticity can give rise to cells exhibiting elevated levels of CD44 across lineages in cancers and healthy tissues. Furthermore, cancer cells of a mesenchymal origin and activated T-cells involved in immune responses can also display CD44 at the plasma membrane. In the context of cancer, diseased mesenchymal cells contribute to drug resistance (Brabletz et al., 2018), and CD44 itself has been shown to confer metastatic potential to these cells (Günthert et al., 1991). However, molecular mechanisms underlying a functional role of CD44 are not fully understood.

CD44 has been proposed to act as a ligand-binding surface protein regulating cell adhesion and migration through the extracellular matrix, and as a membrane co-receptor mediating intracellular signal transduction (Ponta et al., 2003). Importantly, CD44 interacts with and mediates endocytosis of hyaluronates (Hyal) (Aruffo et al., 1990; Hua et al., 1993), a class of glycosaminoglycan biopolymers that can promote EMT and tumor development (Bartolazzi et al., 1994; Toole, 2004; Zoltan-Jones et al., 2003).

Hyal contain negatively-charged carboxylates for every disaccharide unit. This raises the question of what counter ions these biopolymers interact with at physiological pH during endocytosis, to enable the uptake of Hyal without altering the intracellular ionic balance and membrane potentials. The ability of Hyal to interact with iron (Fe) (Mercê et al., 2002), together with the iron dependency of mesenchymal cancer cells *in vivo* and their susceptibility to ferroptotic cell death (Basuli et al., 2017; Hangauer et al., 2017; Mai et al., 2017; Schonberg et al., 2015; Torti and Torti, 2013; Viswanathan et al., 2017), prompted us to search for a putative CD44-mediated iron uptake mechanism.

Here, we report a novel hyaluronate-dependent iron endocytosis pathway mediated by CD44 that is upregulated during EMT. We found that increase of iron uptake is required for the activity of an iron-dependent demethylase regulating the expression of specific genes in the mesenchymal state of cells. This work illuminates a central role of iron in the regulation of epigenetic plasticity, functionally linking the membrane glycoprotein CD44 to the nucleus.

## RESULTS

### CD44 mediates hyaluronate-dependent iron endocytosis

To characterize interactions occurring between Hyal and iron, we employed nuclear magnetic resonance (NMR) spectroscopy. Addition of Fe(III) to a low-molecular-mass Hyal led to line broadening of proton signals indicating that this organic substrate interacts with the metal at neutral pH (Figure 1A). Acidifying the sample to protonate the carboxylates of Hyal released the metal, thereby rescuing the proton signals of free Hyal. This demonstrates the dynamic and reversible nature of metal coordination by Hyal in near-physiological conditions of solvent, temperature and pH.

**Figure 1.**
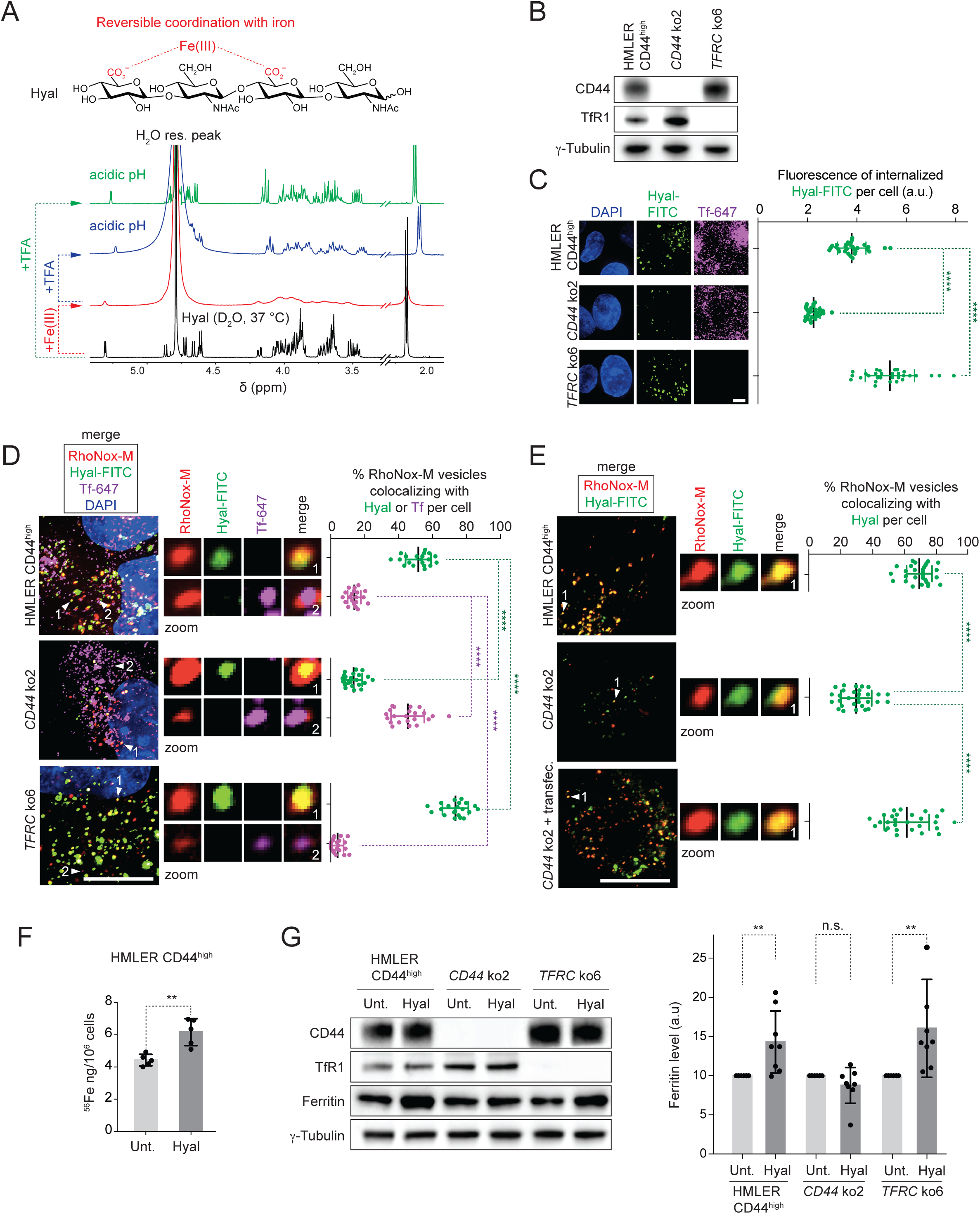
CD44 mediates hyaluronate-dependent iron endocytosis. (**A**) Molecular structure (top) and ^1^H NMR spectra of Hyal (bottom). Data showing that Hyal interacts with Fe(II) in near-physiological conditions. These interactions are reversed upon acidification of the media using trifluoroacetic acid (TFA). Functional groups that can interact with iron are highlighted in red. (**B**–**G**) Parental HMLER CD44^high^ and corresponding ko cell lines were used throughout these panels. (**B**) Western blot analysis of CD44 and TfR1 levels in *CD44* and *TFRC* ko clones generated using CRISPR-Cas9. (**C**) Fluorescence microscopy analysis of internalized Hyal-FITC. Data showing that knocking out *CD44* reduces endocytosis of Hyal. *n* = 3 biological replicates. (**D**) Fluorescence microscopy analysis of RhoNox-M-positive vesicles colocalizing with internalized Hyal-FITC or Tf-647. Data showing that cells expressing CD44 exhibit RhoNox-M/Hyal-positive vesicles that are free of Tf. The proportion of these vesicles is significantly reduced by knocking out *CD44. n* = 3 biological replicates. (**E**) Fluorescence microscopy analysis of RhoNox-M-positive vesicles colocalizing with Hyal-FITC in a *CD44* ko clone complemented with *CD44*. Data showing that *CD44* complementation restores a higher proportion of RhoNox-M/Hyal-positive vesicles. *n* = 3 biological replicates. (**F**) ICP-MS measurements of cellular iron. Data showing that supplementing CD44-expressing cells with Hyal increases iron uptake. *n* = 5 biological replicates. (**G**) Western blot analysis of ferritin levels representative of 8 biological replicates. Data showing that supplementing CD44-expressing cells with Hyal increases levels of the iron storage protein ferritin. DAPI, 4′,6-diamidino-2-phenylindole. Unt., untreated. Scale bars, 10 μm. Bars and error bars, mean values ± s.d. * *P* ≤ 0.05, ** *P* ≤ 0.01, *** *P* ≤ 0.001, **** *P* ≤ 0.0001; n.s., not significant; unpaired *t*-tests. See also Figure S1.

To evaluate the potential of high-molecular-weight Hyal-based organometallic complexes to be internalized by CD44, we knocked out (ko) *CD44* from mesenchymal human mammary HMLER cells stably expressing high levels of CD44 (HMLER CD44^high^) (Figure 1B) (Morel et al., 2008). For comparison, we independently knocked out *TFRC*, the gene coding for transferrin receptor protein 1 (TfR1), which mediates endocytosis of iron-bound transferrin (Tf) and plays a central role in the regulation of cellular iron homeostasis. We then evaluated the capacity of parental cells and ko clones to take up fluorescently labeled Hyal. Endocytosis of Hyal was reduced in *CD44* ko and increased in *TFRC* ko clones compared to parental cells, validating the functional role of CD44 in mediating this process (Figure 1C).

Hyal and the iron-carrier Tf, which normally interacts with TfR1, predominantly localized in distinct vesicles in parental cells (Figure 1D), allowing for a comparative analysis of metal uptake mediated either by CD44 or TfR1. Thus, we set out to evaluate whether acidic vesicles contained Fe(II) together with Hyal independently of Tf. To this end we used the Fe(II)-specific turn-on fluorescent probe RhoNox-M, which can be employed to detect the presence of Fe(II) in lysosomes (Niwa et al., 2014). Parental cells exhibited RhoNox-M/Hyal-positive vesicles that were free of Tf as defined by fluorescence microscopy. *CD44* ko cells exhibited a lower proportion of RhoNox-M/Hyal-positive vesicles, while *TFRC* ko cells showed the opposite trend along with a reduction in RhoNox-M/Tf-positive vesicles compared to parental cells (Figure 1D). Complementing ko cells with *CD44* restored a higher proportion of RhoNox-M/Hyal-positive vesicles (Figure 1E). Importantly, RhoNox-M/Hyal-positive vesicles free of Tf could also be found in other breast cancer cell lines, primary breast cancer cells, circulating tumor cells (CTC) of lung and colon cancers and in primary human T-cells, illustrating the general nature of this pathway (Figures S1A and S1B).

In line with CD44 mediating iron endocytosis, supplementing cells with Hyal increased levels of cellular iron in parental CD44^high^ and *TFRC* ko but not in *CD44* ko cells as defined by the fluorescence of RhoNox-M quantified by flow cytometry (Figure S1C). This was observed across cancer cell lines and primary cells, indicating a general feature of cells expressing CD44 (Figure S1D). In CD44-expressing MDA-MB-468 breast cancer cells, iron uptake was more pronounced upon treatment with higher molecular weight Hyal (Figure S1E). Conversely, treating MDA-MB-468 cells with hyaluronidase or blocking antibodies against CD44 or TfR1 reduced iron uptake (Figures S1F and S1G). Additionally, cellular iron was higher in HMLER CD44^high^ cells supplemented with Hyal as measured by inductively coupled plasma mass spectrometry (ICP-MS) (Figure 1F). Finally, treating HMLER CD44^high^ cells with Hyal also led to increased levels of the iron storage protein ferritin in a CD44-dependent manner, further demonstrating increased iron uptake in these conditions (Figure 1G). Collectively, these data show that CD44 mediates iron endocytosis in a Hyal-dependent manner.

### Prevalence of CD44-mediated iron endocytosis in the mesenchymal state of cells

Mesenchymal states of cells are characterized by elevated levels of CD44 in several lineages (Stuelten et al., 2010). To investigate the cellular changes reliant on CD44 levels, we triggered EMT in triple-negative breast cancer MDA-MB-468, luminal breast cancer MCF7 and HMLER CD44^low^ cell lines, primary breast cancer cells and primary lung CTC using epidermal growth factor (EGF), oncostatin M (OSM) or transforming growth factor beta (TGF-β). Upon EGF treatment, MDA-MB-468 cells shifted from an epithelial to a mesenchymal phenotype according to cell morphology, reduced levels of the epithelial marker E-cadherin and increased levels of CD44 (Figure 2A). Consistent with EMT induction, levels of the EMT transcription factor snail increased along with that of the mesenchymal markers vimentin and fibronectin (Figure 2B). Remarkably, CD44 levels increased together with ferritin, whereas levels of TfR1 were reduced, indicating that iron uptake is predominantly mediated by CD44 in these conditions (Figure 2B). In line with this, CD44 loading at the plasma membrane was enhanced during EMT, which was in contrast to a reduced loading of TfR1 (Figures 2C and S2A). The number of RhoNox-M/Hyal-positive vesicles increased together with cellular iron (Figures 2D, 2E and S2B). Importantly, knocking down CD44 antagonized the production of ferritin in cells treated with EGF (Figure 2F).

**Figure 2.**
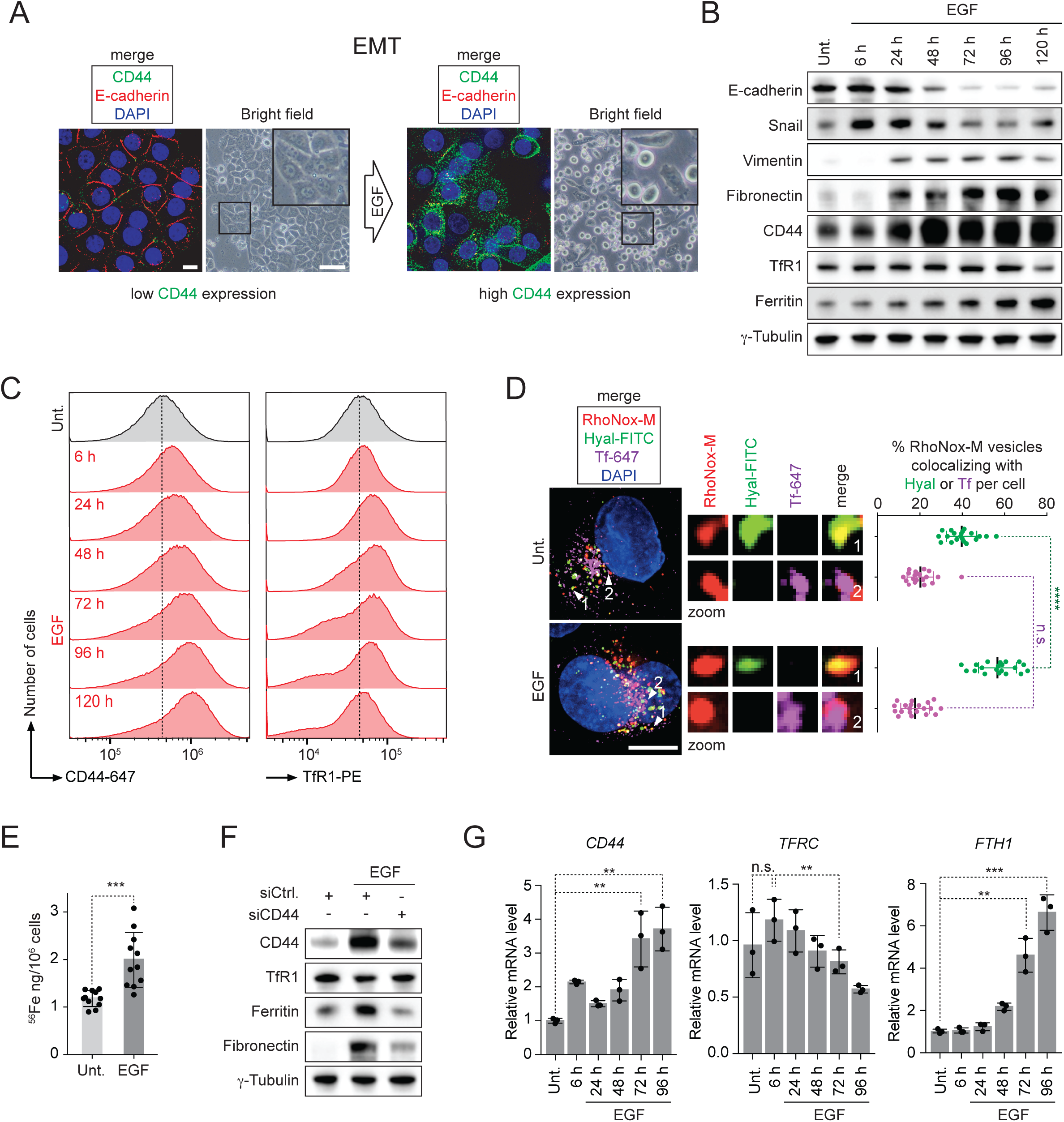
CD44-mediated iron endocytosis prevails in the mesenchymal state of cells. MDA-MB-468 cells were used throughout the figure and were treated with EGF for 72 h, unless stated otherwise. (**A**) Fluorescence microscopy analysis of E-cadherin and CD44 (scale bars, 10 μm). Bright field microscopy analysis of cell morphology (scale bars, 100 μm). Data showing induction of EMT upon EGF treatment. (**B**) Time course western blot analysis of levels of iron homeostasis and EMT proteins. Data showing that, in contrast to TfR1, CD44 levels increase during EMT. (**C**) Time course flow cytometry analysis of CD44 and TfR1 loading at the plasma membrane. Data showing that, in contrast to TfR1, CD44 loading increases steadily during EMT. (**D**) Fluorescence microscopy analysis of RhoNox-M-positive vesicles colocalizing with internalized Hyal-FITC or Tf-647. Data showing that EMT induction is associated with an increased proportion of RhoNox-M/Hyal-positive vesicles. *n* = 3 biological replicates. Scale bar, 10 μm. (**E**) ICP-MS measurements of cellular iron. Data showing that levels of cellular iron increase during EMT. *n* = 11 biological replicates. (**F**) Western blot analysis of ferritin levels in CD44 knock down conditions. Data showing that knocking down CD44 antagonizes the production of the iron storage protein ferritin in cells treated with EGF. (**G**) Time course RT-qPCR analysis of *CD44, TFRC* and *FTH1* mRNA levels. Data indicating that, in contrast to *TFRC*, the RNA transcript of *CD44* increases concomitantly with that of *FTH1* during EMT. *n* = 3 biological replicates. Bars and error bars, mean values ± s.d. * *P* ≤ 0.05, ** *P* ≤ 0.01, *** *P* ≤ 0.001, **** *P* ≤ 0.0001; n.s., not significant; unpaired *t*-tests. See also Figure S2.

Furthermore, RT-qPCR showed that mRNA levels of *CD44*, and *FTH1* coding for ferritin, increased during EMT, whereas that of *TFRC* decreased (Figure 2G). This data was in agreement with the well-established negative feedback loop regulating the biosynthesis of TfR1 at the translational level by excess cytosolic iron (Muckenthaler et al., 2017). It also indicated that in contrast to *TFRC, CD44* is not subjected to this regulatory loop. Together, these results advocate for an alternative glycan-mediated iron endocytosis pathway reliant on CD44 that prevails in the mesenchymal state of cells.

### EGF-induced EMT is characterized by a redox signature implicating iron

To identify iron-dependent cellular functions required during EMT or for the maintenance of mesenchymal states of cells, we employed a combination of quantitative proteomics and metabolomics. Out of 5,574 proteins that were detected by mass spectrometry, 255 proteins were downregulated while 276 were upregulated in MDA-MB-468 cells treated with EGF (Figure 3A). Gene ontology (GO) revealed a bias towards proteins with oxidoreductase activity being increased in expression in the mesenchymal state (Figure S3A and Table S1). The response to EGF treatment was further characterized by changes in EMT markers, proteins linked to metastasis, proteins involved in the regulation of endocytosis and iron homeostasis as well as metabolic enzymes and an iron-dependent demethylase. For example, EMT was characterized by reduced levels of the epithelial proteins BCAM and CADM-4, whereas vimentin was upregulated. The pro-metastatic protein CD109 was upregulated (Chuang et al., 2017), which was in agreement with the notion that mesenchymal cancer cells are capable of dissemination. Consistent with increased iron uptake during EMT, levels of ferritin increased together with sorting nexin 9 (SNX9), a protein that regulates endocytosis of CD44, which has also been implicated in metastasis (Bendris et al., 2016). Conversely, levels of Tf were reduced, further supporting the prevalent functional role of CD44 in mediating iron endocytosis during EMT. Importantly, we detected an increase of the iron-dependent histone demethylase PHF8, which is involved in active transcription, brain development, cell cycle progression as well as the regulation of EMT *in vivo* and in breast tumor growth (Feng et al., 2010; Fortschegger et al., 2010; Liu et al., 2010; Qi et al., 2010; Shao et al., 2017). In contrast, levels of other demethylases that could be detected by mass spectrometry did not change significantly.

**Figure 3.**
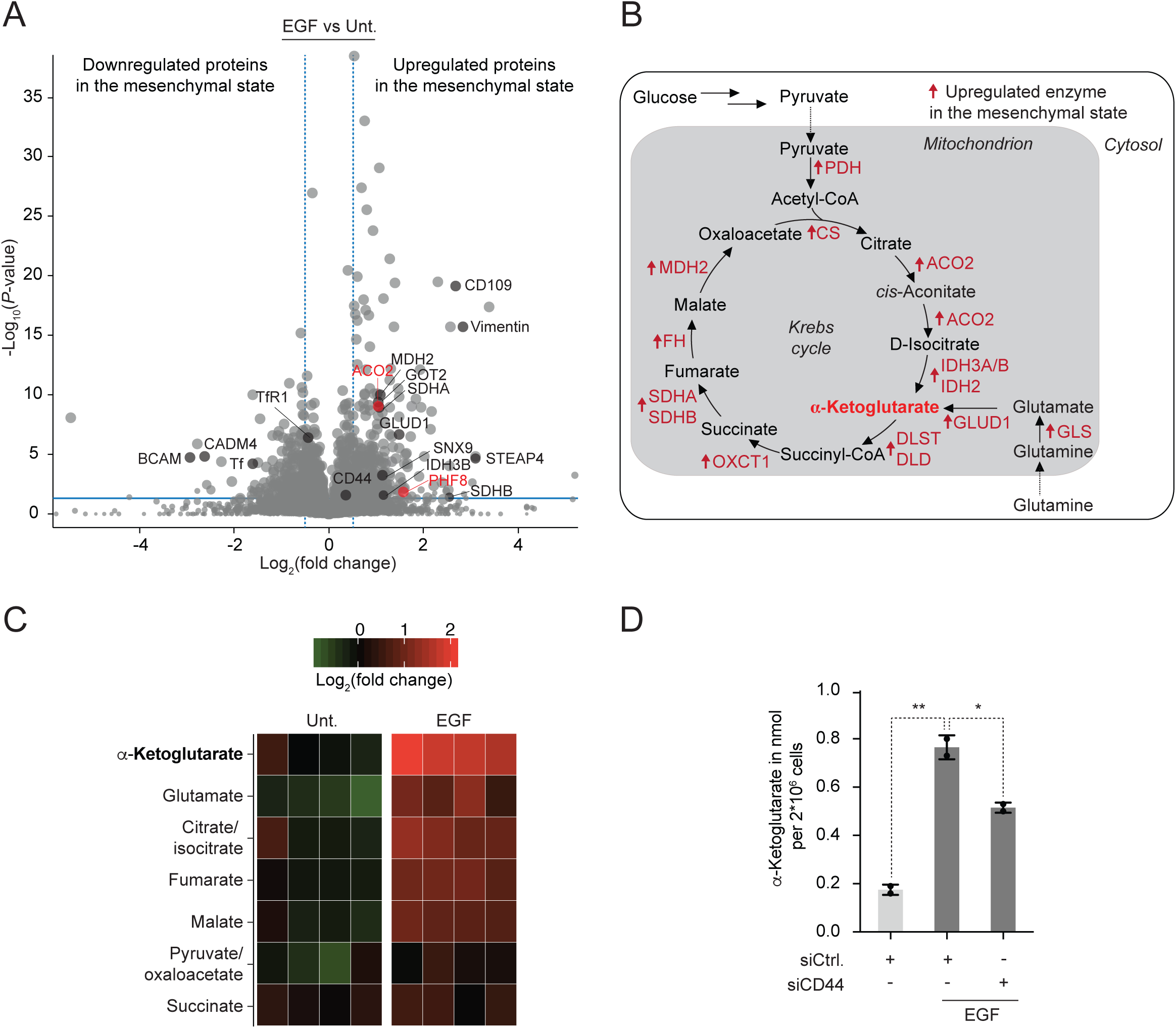
EGF-induced EMT is characterized by a redox signature implicating iron. MDA-MB-468 cells were used throughout the figure and were treated with EGF for 72 h, unless stated otherwise. (**A**) Quantitative label-free proteomics. Volcano plot representing protein expression fold changes. Upregulated (right) and downregulated (left) proteins are shown. Dashed blue line defines a fold change of 1.42, plain blue line defines a *P*-value of 0.05. Data showing that levels of the mitochondrial iron-sulfur cluster-containing protein ACO2 (red) and that of the nuclear iron-dependent demethylase PHF8 (red) increase during EMT. *n* = 3 biological replicates. (**B**) Schematic illustration of metabolic pathways involved in the production of αKG, highlighting enzymes that are upregulated at the protein level during EMT (red). (**C**) Quantitative metabolomics. Heatmap of upregulated (red) and downregulated (green) metabolites in cells treated with EGF for 60 h. Data showing that levels of αKG increase during EMT. *n* = 4 technical replicates. (**D**) Quantification of αKG levels in CD44 knock down conditions using an αKG titration assay. Data showing that knocking down CD44 antagonizes the production of αKG in cells treated with EGF. *n* = 2 technical replicates. Bars and error bars, mean values ± s.d. * *P* ≤ 0.05, ** *P* ≤ 0.01, *** *P* ≤ 0.001, **** *P* ≤ 0.0001; n.s., not significant; unpaired *t*-tests. See also Figure S3.

Metabolic enzymes involved in the conversion of glutamate and pyruvate to α-ketoglutarate (αKG), including glutamate dehydrogenase 1 (GLUD1), isocitrate dehydrogenase 2 (IDH2) and the iron-sulfur cluster-containing aconitase 2 (ACO2), were upregulated along with other enzymes of the Krebs cycle (Figures 3A and 3B). αKG is a co-substrate required for the iron-catalyzed oxidative demethylation of histone and nucleic acid methyl marks, and is implicated in the maintenance of pluripotency (Carey et al., 2015). Consistent with the upregulation of these mitochondrial enzymes, quantitative metabolomics indicated a marked increase of αKG and its precursors glutamate and isocitrate in MDA-MB-468 cells treated with EGF (Figures 3C and S3B, Table S2). We then interrogated whether reducing iron uptake could potentially alter mitochondrial metabolism in cells treated with EGF. Knocking down CD44 antagonized the upregulation of αKG in cells treated with EGF (Figure 3D), which was consistent with a role of iron in mitochondria during EMT. These biochemical changes we observed during EMT, including enhanced cellular uptake of iron together with the upregulation of PHF8 and its co-substrate αKG, further hinted toward a specific regulation occurring in the nucleus at the chromatin level (Li et al., 2018).

### Nuclear iron is a rate-limiting regulator of epigenetic plasticity

Metals are ubiquitous in cell biology, acting as protein cofactors. Due to its unique electron configuration, iron distinguishes itself by its capacity to catalyze oxidative demethylation of protein residues, including those of histones, and methylated nucleobases. Thus, iron is a rate-limiting factor of these processes (Figure 4A). In line with a role of iron in the nucleus of cells undergoing EMT, subcellular fractionation indicated higher levels of nuclear ferritin in MDA-MB-468 cells undergoing EMT (Figure 4B) (Smith et al., 1990).

**Figure 4.**
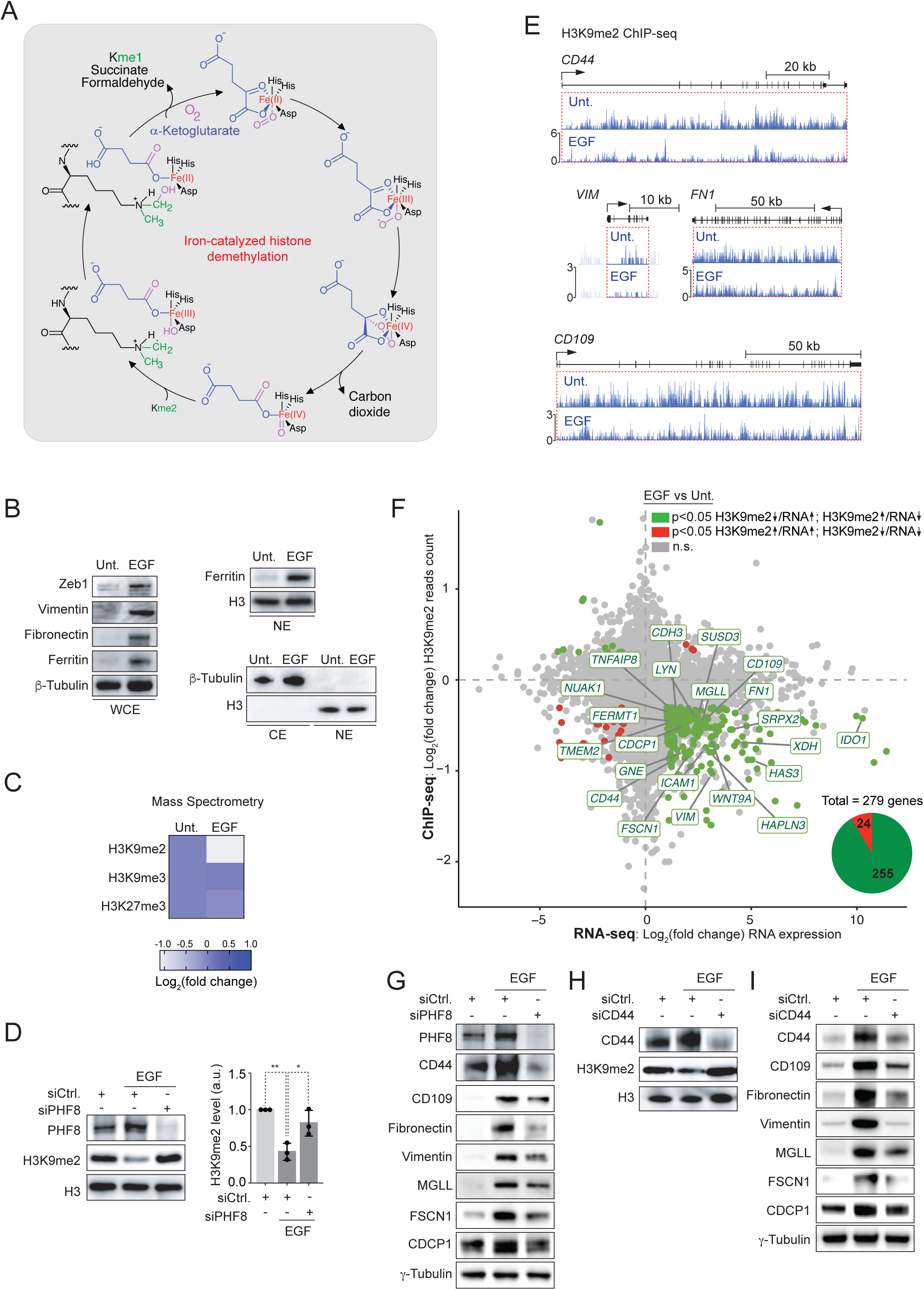
Nuclear iron regulates epigenetic plasticity through oxidative histone demethylation. (**A**) Schematic illustration of the catalytic cycle of iron-mediated oxidative demethylation of a methylated lysine residue. Active site of PHF8 enzyme (black) bound to iron (red), together with molecular oxygen (pink) and αKG (blue) co-substrates required for demethylation are shown. (**B**–**I**) MDA-MB-468 cells were used throughout the figure and were treated with EGF for 72 h. (**B**) Subcellular fractionation and western blot analysis of nuclear ferritin levels. NE, nuclear extract; CE, cytoplasmic extract; WCE, whole cell extract. Data showing increased levels of the iron storage protein ferritin in the nucleus of mesenchymal cells. (**C**) Quantitative mass spectrometry. Heatmap of H3K9me2, H3K9me3 and H3K27me3 levels showing a reduction of the repressive histone mark H3K9me2 in the mesenchymal state of cells. *n* = 5 biological replicates. *P* ≤ 0.0001 for H3K9me2 and n.s. for H3K9me3 and H3K27me3; two-sided associated *t*-tests. (**D**) Western blot analysis of levels of H3K9me2 in PHF8 knock down conditions. Data showing that knocking down PHF8 antagonizes the reduction of H3K9me2 induced by EGF. Bars and error bars, mean values ± s.d. * *P* ≤ 0.05, ** *P* ≤ 0.01, *** *P* ≤ 0.001, **** *P* ≤ 0.0001; n.s., not significant; unpaired *t*-tests. (**E**) H3K9me2 ChIP-seq profiles of selected genes. Data showing the reduction of H3K9me2 ChIP-seq reads count in the gene body of these mesenchymal and pro-metastatic genes. (**F**) Scatter plot correlation of H3K9me2 ChIP-seq reads count in genes (*n* = 3 biological replicates) and RNA-seq (*n* = 2 biological replicates). Data showing that reduction of H3K9me2 reads count in specific genes correlates with increased expression of these genes. (**G**) Western blot analysis of proteins whose genes are regulated by H3K9me2 in PHF8 knock down conditions. Data showing that knocking down PHF8 antagonizes the upregulation of iron-regulated genes in cells treated with EGF. (**H**) Western blot analysis of H3K9me2 levels in CD44 knock down conditions. Data showing that knocking down CD44 antagonizes the reduction of H3K9me2 in cells treated with EGF. (**I**) Western blot analysis of levels of proteins whose genes are regulated by H3K9me2 in CD44 knock down conditions. Data showing that knocking down CD44 antagonizes the upregulation of iron-regulated genes in cells treated with EGF. See also Figure S4.

The iron- and αKG-dependent demethylase PHF8 has been shown to promote demethylation of H3K9me2 (Yu et al., 2010), a repressive histone mark whose reduction has been implicated in solid tumors (McDonald et al., 2017; McDonald et al., 2011; Zhao et al., 2016). Increase of PHF8 in MDA-MB-468 cells undergoing EMT correlated with the preferential reduction of H3K9me2 compared to the other repressive histone marks H3K9me3 and H3K27me3, as defined by quantitative mass spectrometry (Figure 4C and Table S3). Additionally, knocking down PHF8 prevented demethylation of H3K9me2 in MDA-MB-468 cells treated with EGF (Figure 4D), validating a functional role of this demethylase during EMT.

Next, we examined the genome-wide distribution of H3K9me2 by chromatin immunoprecipitation sequencing (ChIP-seq) to identify PHF8-regulated genes whose expressions are iron-dependent. ChIP-seq revealed large organized heterochromatin K9 modifications, previously termed LOCKs (McDonald et al., 2017; McDonald et al., 2011). These data showed a reduction of H3K9me2 ChIP-seq reads count in the body of specific genes in cells treated with EGF, including *CD44, CD109, VIM* and *FN1* coding for vimentin and fibronectin, respectively (Figure 4E and Table S4). These data indicate that CD44 positively regulates its own expression at the transcriptional level by mediating iron endocytosis, unlike TfR1 that is positively regulated at the translational level *via* iron responsive elements (IRE) under iron-depleted conditions (Muckenthaler et al., 2017). Furthermore, analysis of the transcriptome by RNA-seq showed that a reduction of H3K9me2 reads count correlated with increased gene expression in agreement with the repressive nature of H3K9me2 (Figure 4F and Table S4). For instance, out of 279 genes where reduction of H3K9me2 was observed, 255 genes were transcriptionally upregulated. In addition to *CD44*, we identified genes involved in cancer, regulation of EMT, development, immune responses, inflammation, and wound healing, which are all processes CD44 is linked to (Figure S4). In comparison, we did not observe a significant reduction of H3K9me2 in genes coding for EMT transcription factors. However, we identified long non-coding (lnc) RNA genes previously reported to be involved in cancer (Figure S4). In agreement with PHF8 playing a crucial role in regulating these genes through oxidative demethylation of H3K9me2, knocking down this demethylase partly blocked EGF-induced expression of iron-regulated genes, including CD44 (Figure 4G). Remarkably, knocking down CD44 phenocopied PHF8 knock down in MDA-MB-468 cells, partly blocking H3K9me2 demethylation and the expression of genes regulated by PHF8 (Figures 4H and 4I). Taken together, these data establish a direct functional link between CD44 and the regulation of gene expression involving iron endocytosis and iron-catalyzed histone demethylation.

### Targeting iron homeostasis interferes with the maintenance of a mesenchymal state of cells

Small molecules provide a means to manipulate cellular processes with valuable spatial and temporal resolution (Gerry and Schreiber, 2018). To directly address the implication of the nuclear iron pool, we set out to identify small molecule chelators of iron that could specifically target the nucleus. To this end, we functionalized known iron chelators with alkyne functional groups that can be chemically labeled in cells by means of click chemistry (Cañeque et al., 2018). Fluorescent labeling of a clickable surrogate of deferoxamine (cDFO) in cells revealed a predominant nuclear localization of this small molecule (Figure 5A). Thus, deferoxamine (DFO) represents a suitable tool to directly interrogate functional roles of iron in the nucleus. For instance, DFO prevented the demethylation of H3K9me2 in cancer cell lines and primary cancer cells that were treated either with EGF, OSM or TGF-β (Figures 5B and S5A). Consistently, DFO antagonized the upregulation of proteins whose genes were found to be regulated by PHF8 in these conditions (Figures 5C and S5B). These data provide direct evidence of the functional role of nuclear iron in the regulation of the expression of these genes.

**Figure 5.**
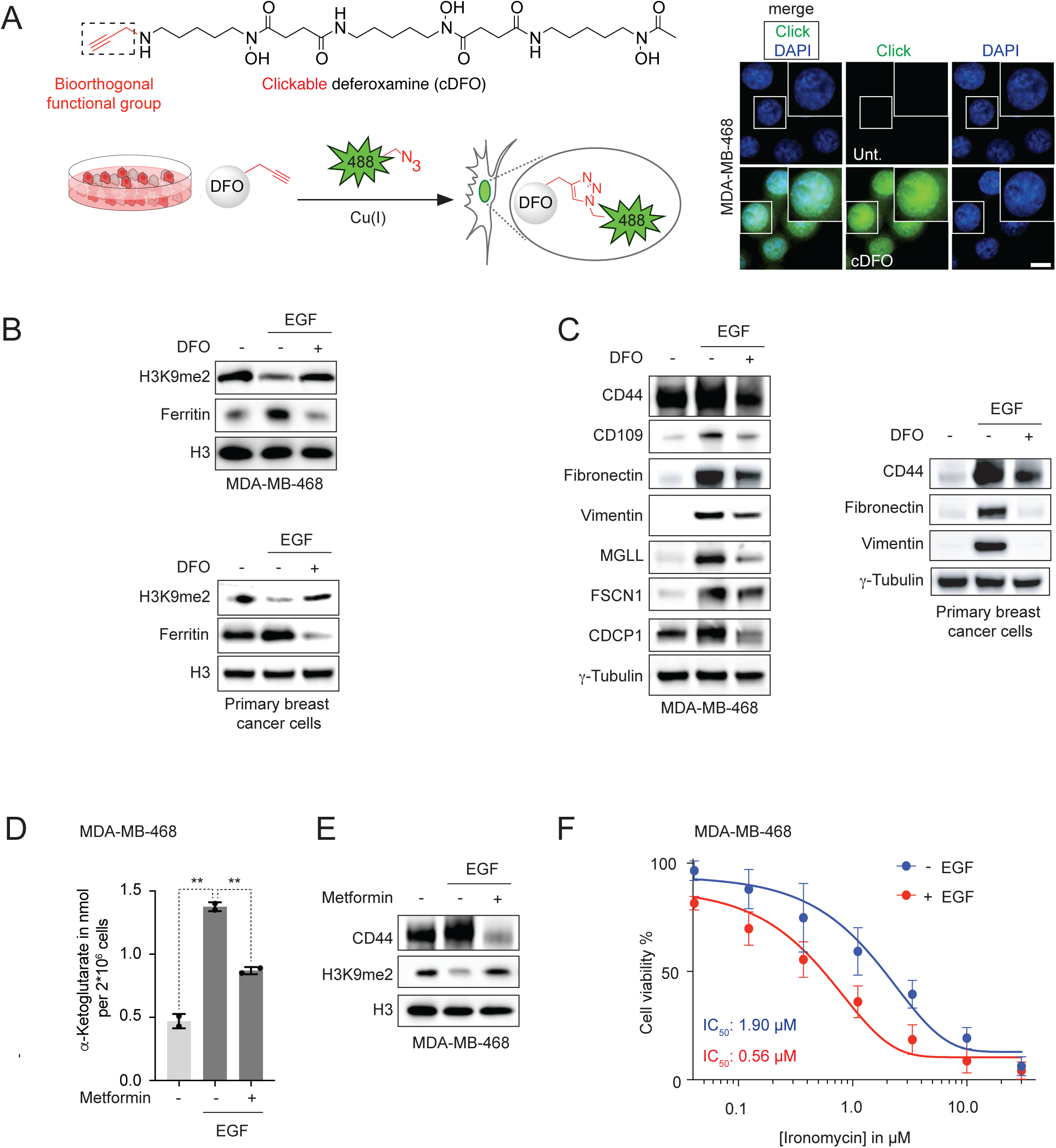
Targeting of iron-regulated processes. (**A**) Molecular structure of cDFO (top), schematic illustration of the chemical labeling of cDFO in cells (bottom) and corresponding fluorescence microscopy images (right). 488 represents the Alexa-Fluor-488 fluorophore and has been chosen arbitrarily. Data showing that labeled DFO colocalizes with DAPI, preferentially targeting the nucleus. Scale bar, 10 μm. (**B**) Western blot analysis of H3K9me2 levels in cells cotreated with EGF and DFO. Data showing that targeting nuclear iron antagonizes the effect of EGF, preventing the reduction of H3K9me2. (**C**) Western blot analysis of proteins whose genes are regulated by H3K9me2 in cells cotreated with EGF and DFO. Data showing that targeting nuclear iron antagonizes the effect of EGF, preventing the upregulation of iron-regulated genes. (**D**) Quantification of αKG using an αKG titration assay (*n* = 2 technical replicates) and western blot analysis of H3K9me2 levels in cells cotreated with EGF and metformin. Data showing that targeting mitochondrial metabolism antagonizes the effect of EGF, preventing the reduction of H3K9me2. (**E**) Cell viability curves of cells cotreated with EGF and ironomycin. Data showing that cells in the mesenchymal state are more vulnerable to lysosomal iron targeting compared to cells in the epithelial state. *n* = 3 biological replicates. Bars and error bars, mean values ± s.d. * *P* ≤ 0.05, ** *P* ≤ 0.01, *** *P* ≤ 0.001, **** *P* ≤ 0.0001; n.s., not significant; unpaired *t*-tests. See also Figure S5.

Metformin has been shown to target metals in the mitochondria and to inhibit the Krebs cycle (Janzer et al., 2014; Müller et al., 2018). Consistent with αKG being required for iron-dependent demethylation of H3K9me2 in the nucleus, metformin antagonized EGF-induced upregulation of αKG (Figure 5D), partly blocking H3K9me2 demethylation and CD44 expression (Figure 5E). In addition, a phosphonium derivative of DFO (pDFO) designed to selectively target mitochondrial iron, interfered with αKG production in EGF treated cells, antagonizing H3K9me2 demethylation and CD44 expression (Figure S5C). These data validate a functional role of mitochondrial iron during EMT and showed that targeting mitochondrial metabolism provides another means to control epigenetic plasticity.

Finally, the lysosomal iron-sequestering small molecule ironomycin has previously been shown to effectively kill mesenchymal and tumor-initiating cells *in vitro* and *in vivo* due to a higher iron content in these cells (Mai et al., 2017). Consistently, EGF potentiated the cytotoxic effect of ironomycin against MDA-MB-468 cells, which was in line with the observed increase of iron endocytosis in these conditions, providing a powerful strategy to eradicate tumorigenic cells undergoing EMT (Figure 5F). Together, these data support a mechanism whereby iron regulates epigenetic plasticity (Flavahan et al., 2017), a general pathway prone to small molecule intervention with topological resolution.

## DISCUSSION

This investigation illuminates a unifying mechanism involving CD44, Hyal and iron in the regulation of epigenetic plasticity (Figure 6). Our findings link iron endocytosis to enhanced mitochondrial metabolism and molecular changes occurring at the chromatin level. The free carboxylates of Hyal are prone to interact with iron outside of cells at physiological pH, thereby enabling CD44 to mediate endocytosis of these organometallic complexes. Upon acidification of endocytic vesicles and ensuing protonation of Hyal, iron is released and translocates into the cytosol to traffic towards specific cellular compartments. In particular, upregulation of the iron-dependent demethylase PHF8 and increased levels of nuclear ferritin together with the reduction of the repressive histone mark H3K9me2 in an iron-dependent manner provide solid evidences for a functional role of nuclear iron during EMT.

**Figure 6.**
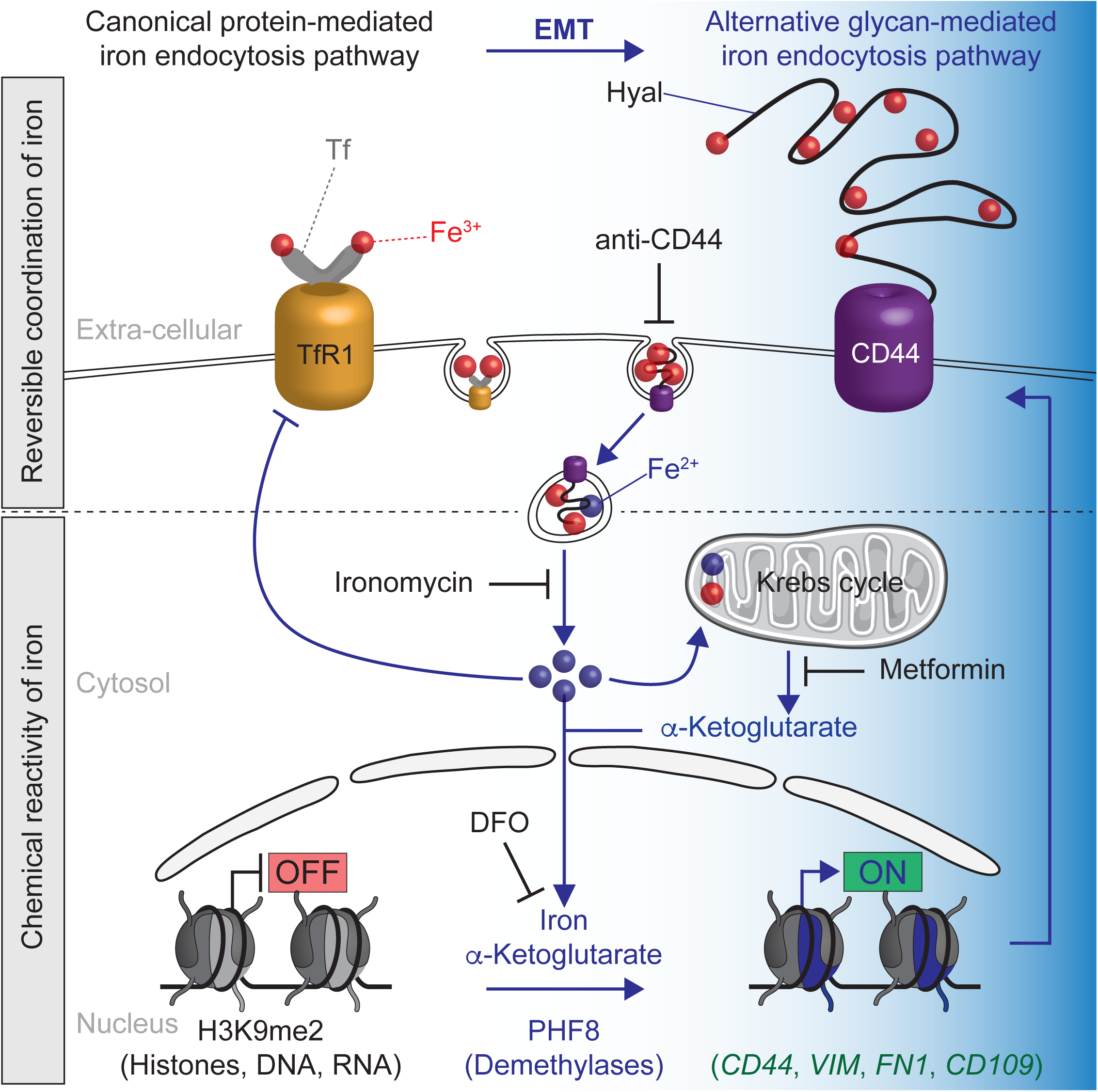
Reciprocal endocytic-epigenetic regulation involving iron. Iron bound to Tf or Hyal enters cells by means of TfR1-or CD44-mediated endocytosis, respectively. Excess cellular iron inhibits the canonical TfR1 pathway. Nuclear iron, αKG produced in mitochondria and iron-dependent enzymes mediate oxidative demethylation of chromatin marks to unlock the expression of specific genes including *CD44*. CD44 regulates its own expression at the transcriptional level by mediating iron endocytosis and this pathway prevails in the mesenchymal state of cells. Iron homeostasis can be targeted at the plasma membrane, the endosomal/lysosomal compartment, the mitochondria and the nucleus using specific antibodies or small molecules.

In this context, two independent iron endocytosis pathways reliant on the protein Tf or the glycan Hyal coexist. Interestingly, these biomolecules represent distinct classes of organic metal carriers of different biosynthetic origins. The CD44-dependent pathway involves Hyal and is switched on at the onset of EMT progressively taking over as the prevalent mechanism of iron endocytosis compared to the TfR1-dependent pathway. Importantly, *CD44* is an iron-regulated gene, indicating a positive feedback loop whereby CD44 regulates its own expression (Figure 6). In contrast, TfR1 is negatively regulated at the translational level by excess iron. This provides a rationale underlying the global increase of CD44 and the reduction of TfR1 during the course of EMT. In epithelial cells, the iron-demand is moderate making the self-regulated TfR1-dependent pathway sufficient to maintain a basal level of iron. However, the higher needs of iron in the mesenchymal state of cells to unlock the expression of mesenchymal genes cannot be solely fulfilled by this pathway. Thus, upregulation of CD44 triggered by growth factors, cytokines and other signaling molecules represent a powerful alternative to increase the cellular iron load.

In the nucleus, using αKG and taking advantage of demethylases including PHF8, iron operates as a metal catalyst mediating oxidative demethylation of histone residues. Iron is therefore a rate-limiting regulator of the expression of specific genes (Figure 6). Hence, to unlock these genes, cells concomitantly upregulate the production of demethylases and increase iron uptake. The repressive histone mark H3K9me2, which is a direct substrate of iron and PHF8, was identified as a key post-translational modification regulating the expression of mesenchymal genes. However, other enzymes involved in methylation, acetylation and deacetylation are expected to work in concert with this iron-dependent demethylation of H3K9me2. Our analysis revealed that H3K9me2 governs the expression of genes involved in cancer, development, immune responses, inflammation, and wound healing. Interestingly, CD44 has previously been linked to these processes, suggesting a functional regulatory role of iron.

Remarkably, cellular iron homeostasis can be altered at the plasma membrane by interfering with iron endocytosis using specific antibodies, or alternatively by controlling the chemical reactivity of this metal in selected cellular compartments using appropriate small molecules (Figure 6). These pharmacological tools provide a means to dissect the processes reliant on iron in various settings and could be further developed for therapeutic intervention.

Intriguingly, high-molecular-mass Hyal confer cancer resistance to naked mole rats and these Hyal have been shown to be refractory to endocytosis due to their larger size (Tian et al., 2013). It is conceivable that such large biopolymers prohibit EMT by sequestering iron outside of cells. Furthermore, while we have shown that Hyal can reversibly interact with iron, other metal ions can potentially be endocytosed using a similar glycosaminoglycan-mediated pathway and may also contribute to the regulation of epigenetic plasticity at different levels. While our study illustrates a functional role of CD44, Hyal and iron in the context of tumorigenic cells, we anticipate that other physiological and pathological processes that rely on the status of distinct histone marks or modified DNA and RNAs, are under the control of similar mechanisms involving iron-dependent demethylases (Gerken et al., 2007; Greer and Shi, 2012; Jia et al., 2011; Tsai et al., 2014).

## SUPPLEMENTAL INFORMATION

Supplemental information includes 5 Figures and 4 Tables and can be found with this article online.

## ACKNOWLEDGMENTS

The R.R. research group is funded by the European Research Council (ERC) under the European Union’s Horizon 2020 research and innovation programme (grant agreement No [647973]), the Fondation Charles Defforey-Institut de France and Ligue Contre le Cancer (Equipe Labellisée). R.R. and D.L. are supported by Region IdF for NMR and MS infrastructures. We thank the PICT-IBiSA@Pasteur Imaging Facility of Institut Curie, member of the France-BioImaging national research infrastructure for the use of microscopes, Christine Gaillet for assistance with NMR spectroscopy, the ICP-MS platform at the Institut de Physique du Globe de Paris, which is supported by IPGP multidisciplinary program PARI and Paris–Region IdF (SESAME grant agreement No [12015908]), Guillaume Arras for assistance with mass spectrometry data analysis, Sylvère Durand and Guido Kroemer for providing access to the metabolomics platform and Patricia Legoix for NGS sample preparation. High-throughput sequencing was performed by the ICGex NGS platform of Institut Curie, supported by ANR-10-EQPX-03 (Equipex) and ANR-10-INBS-09-08 (France Génomique Consortium) from the Agence Nationale de la Recherche (Investissements d’Avenir program), by the Cancéropole IdF and the SiRIC-Curie program – SiRIC Grant (INCa-DGOS-4654). We thank Claire Hivroz and Stéphanie Dogniaux for providing us with primary human T-cells.

## AUTHOR CONTRIBUTIONS

R.R. conceptualized the study and directed the research. R.R., S.M. and F.S. designed the experiments. T.C. performed NMR spectroscopy and synthesized cDFO and pDFO. A.V. synthesized the iron probe RhoNox-M. S.M. produced knock out cell lines and performed the experiments in relation to iron endocytosis including western blotting, cell imaging, RNA interference, flow cytometry and ICP-MS. A.L. performed RT-qPCR and subcellular fractionation experiments. F.S. prepared the samples for quantitative proteomics and metabolomics. B.L. and D.L. carried out quantitative proteomics. F.S. performed next generation sequencing with assistance from A.D., C.V. and S.B. N.S. performed bioinformatics analysis. R.R., S.M., and F.S. analyzed the data and wrote the article.

## METHODS OUTLINE

Detailed methods are provided in the online version of this paper and include the following:

- KEY RESOURCES TABLE
- LEAD CONTACT AND MATERIALS AVAILABILITY
- EXPERIMENTAL MODEL AND SUBJECT DETAILS
  - Cell culture
  - Genome editing
- METHODS DETAILS
  - Reagent conditions
  - Antibody conditions
  - Chemical synthesis and NMR spectroscopy
  - Western blotting
  - Fluorescence microscopy
  - Transfection
  - Flow cytometry
  - Inductively coupled plasma mass spectrometry
  - RNA interference
  - RT-qPCR
  - Proteomics
  - Metabolomics
  - Titration of α-ketoglutarate
  - Subcellular fractionation
  - Histone extraction
  - Quantitative mass spectrometry by parallel-reaction monitoring
  - ChIP-seq
  - ChIP-qPCR
  - RNA-seq
  - ChIP-seq data analysis
  - RNA-seq data analysis
  - Correlative analysis of ChIP-seq and RNA-seq data
  - Chemical labeling of cDFO in cells
  - Cell viability
- QUANTIFICATION AND STATISTICAL ANALYSIS
- DATA AND CODES AVAILABILITY

---

**Figure S1.**
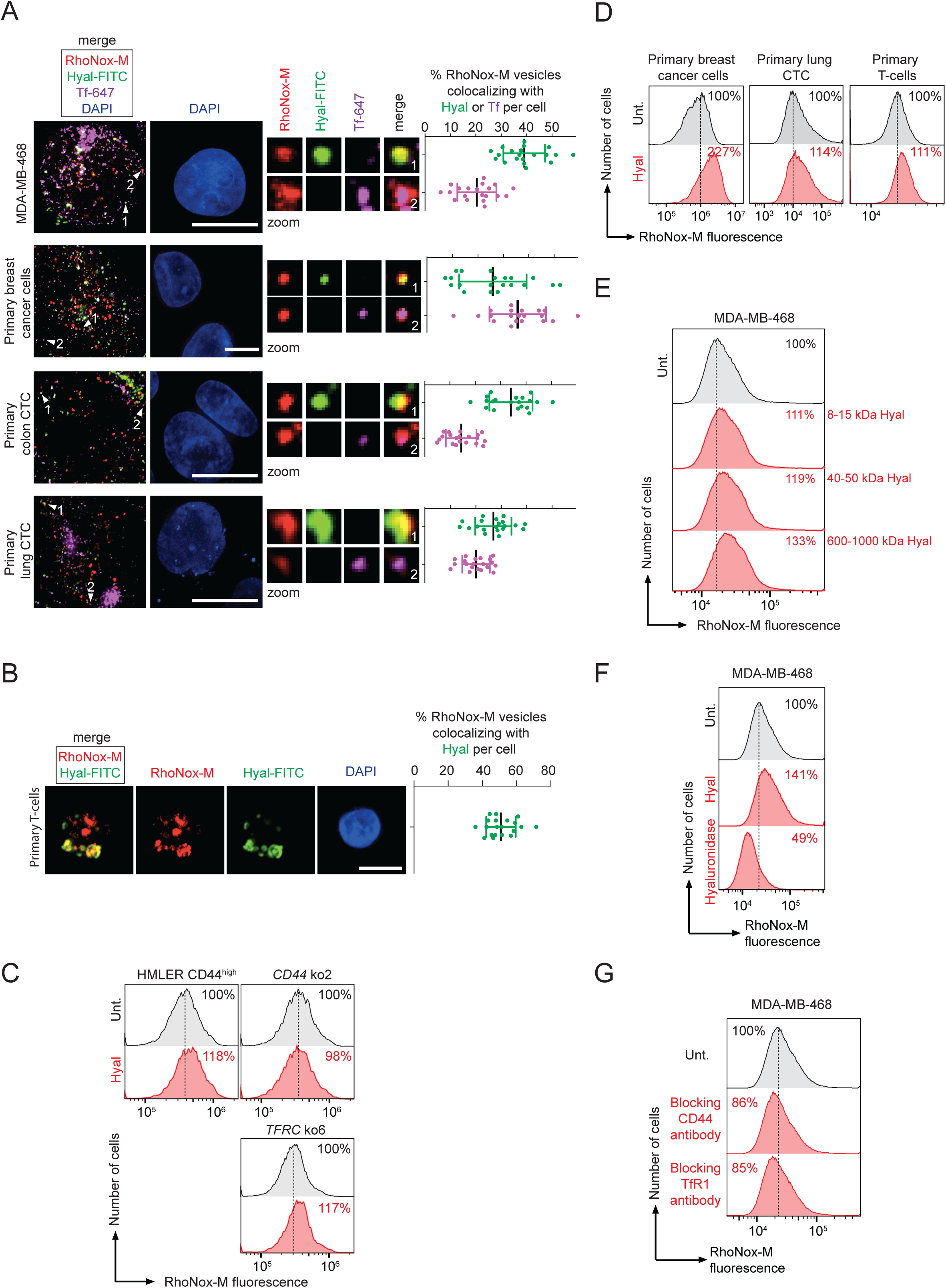
Hyaluronan-dependent iron endocytosis. Related to Figure 1. (**A**) Fluorescence microscopy analysis of RhoNox-M-positive vesicles colocalizing with internalized Hyal-FITC or Tf-647. Data showing that CD44-expressing cells exhibit RhoNox-M/Hyal-positive vesicles free of Tf. *n* = 3 biological replicates for MDA-MB-468 and *n* = 1 for primary cancer cells. Scale bar, 10 μm. (**B**) Fluorescence microscopy analysis of RhoNox-M vesicles colocalizing with internalized Hyal-FITC. Data showing that primary T-cells expressing CD44 exhibit RhoNox-M/Hyal-positive vesicles. No signal of a labeled Tf was detected by fluorescence microscopy in these cells. *n* = 3 biological replicates. Scale bar, 2 μm. Bars and error bars, mean values ± s.d. (**C**) Flow cytometry analysis of RhoNox-M fluorescence in HMLER *CD44* and *TFRC* ko clones. Data showing that supplementing cells with Hyal increases levels of cellular iron in a CD44-dependent manner. (**D**) Flow cytometry analysis of RhoNox-M fluorescence in primary cancer cells and primary T-cells. Data showing that supplementing CD44-expressing cells with Hyal increase levels of cellular iron. (**E**–**G**) Flow cytometry analysis of RhoNox-M fluorescence in cells treated with Hyal, hyaluronidase or blocking antibodies. Data showing that Hyal and CD44 positively regulate levels of cellular iron.

**Figure S2.**
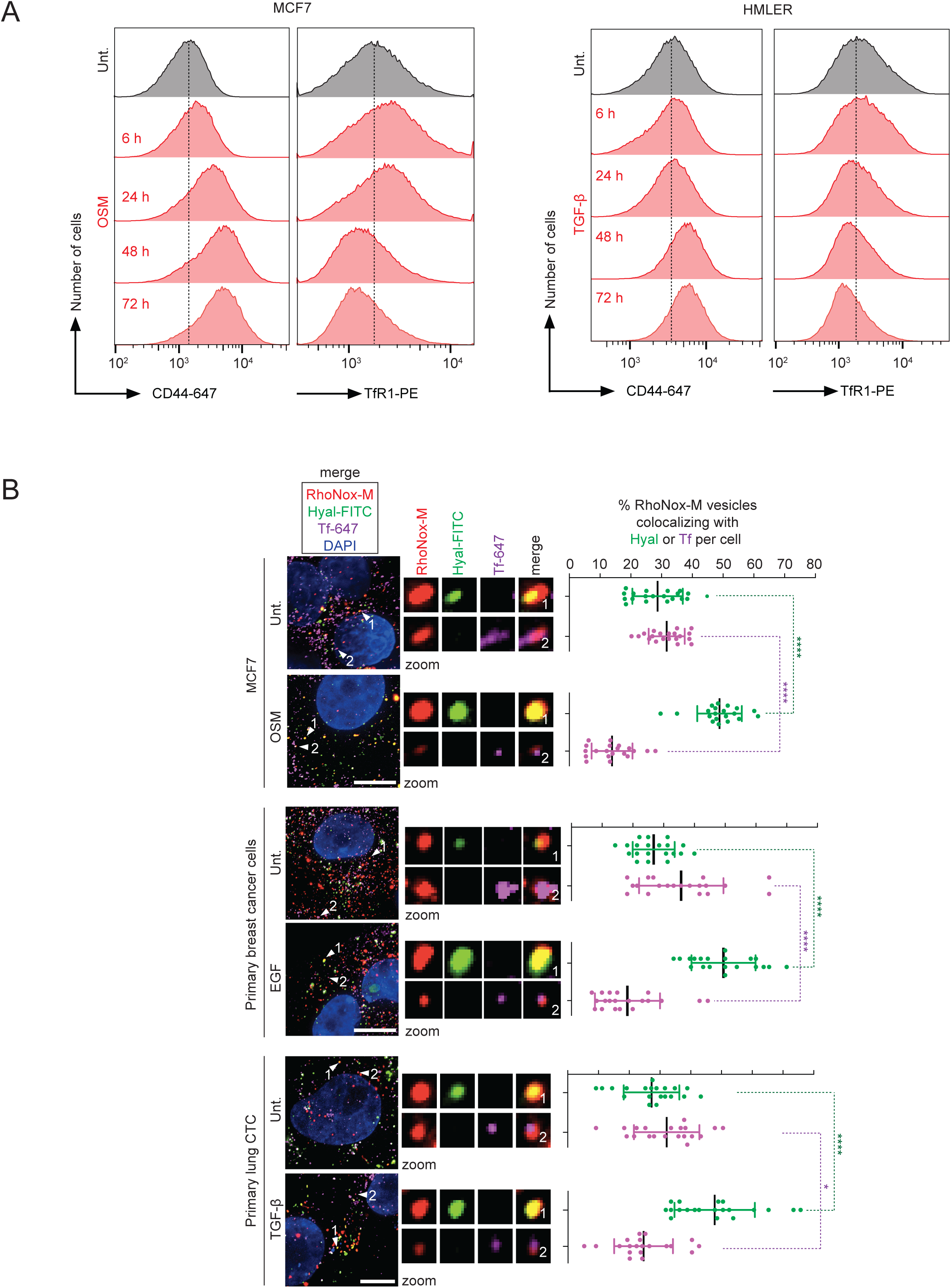
CD44-mediated iron endocytosis prevails in the mesenchymal state of cells. Related to Figure 2. (**A**) Time course flow cytometry analysis of CD44 and TfR1 levels at the plasma membrane. Data showing that, in contrast to TfR1, CD44 loading increases steadily during EMT. (**B**) Fluorescence microscopy analysis of RhoNox-M vesicles colocalizing with internalized Hyal-FITC or Tf-647. Data showing that EMT induction is characterized by an increased proportion of RhoNox-M/Hyal-positive vesicles. *n* = 3 biological replicates for MCF7 cells and *n* = 1 for primary cells. Scale bars, 10 μm. Bars and error bars, mean values ± s.d. * *P* ≤ 0.05, ** *P* ≤ 0.01, *** *P* ≤ 0.001, **** *P* ≤ 0.0001; n.s., not significant; unpaired *t*-tests.

**Figure S3.**
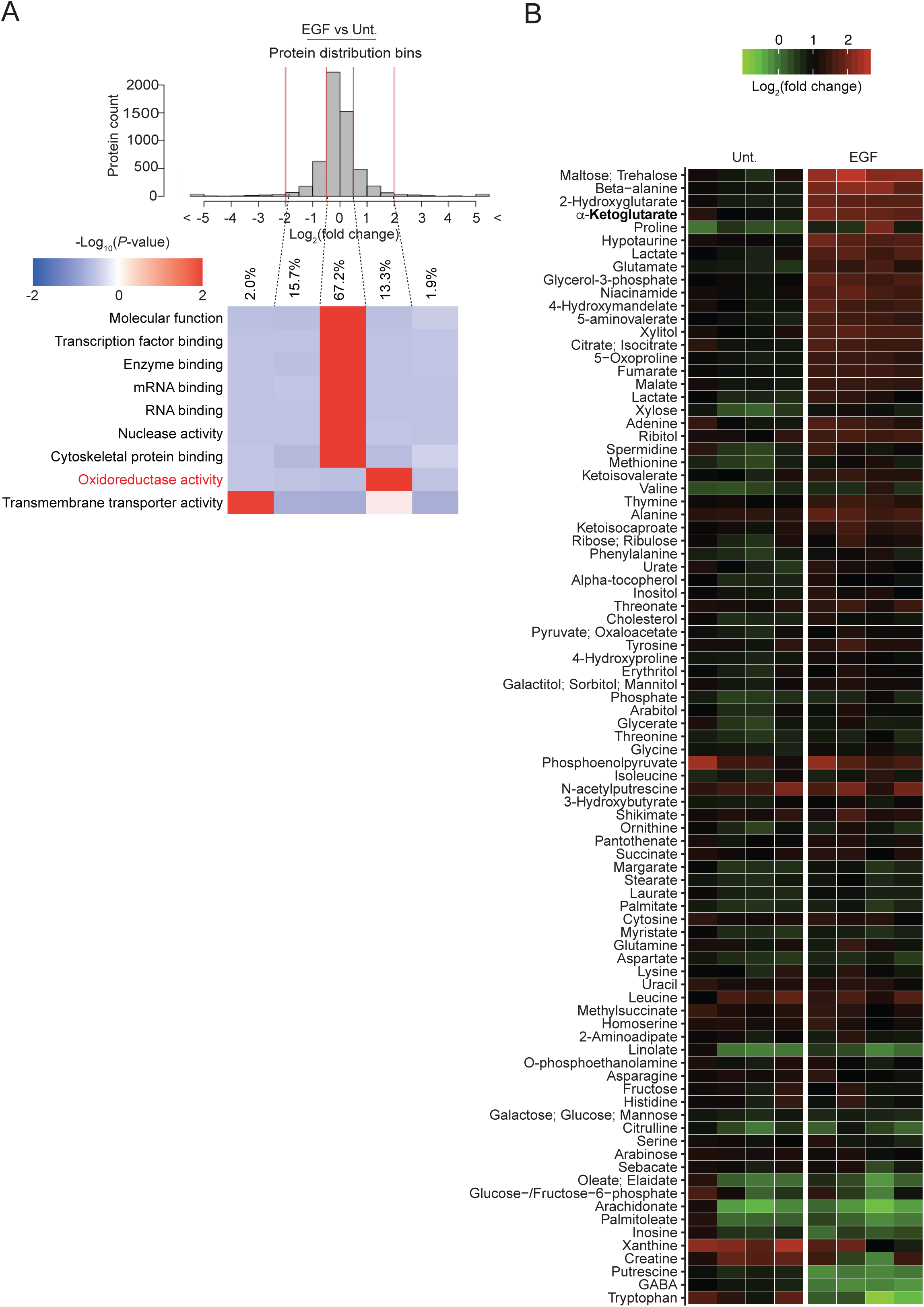
Quantitative Gene Ontology (GO)-term enrichment analysis and metabolome heatmap in cells undergoing EMT. Related to Figure 3. (**A**) Heatmap of proteins ranked by molecular functions illustrating a bias towards proteins with oxidoreductase activity. (**B**) Heatmap of metabolites in cells treated with EGF for 60 h illustrating the increase of specific Krebs cycle metabolites. *n* = 4 technical replicates.

**Figure S4.**
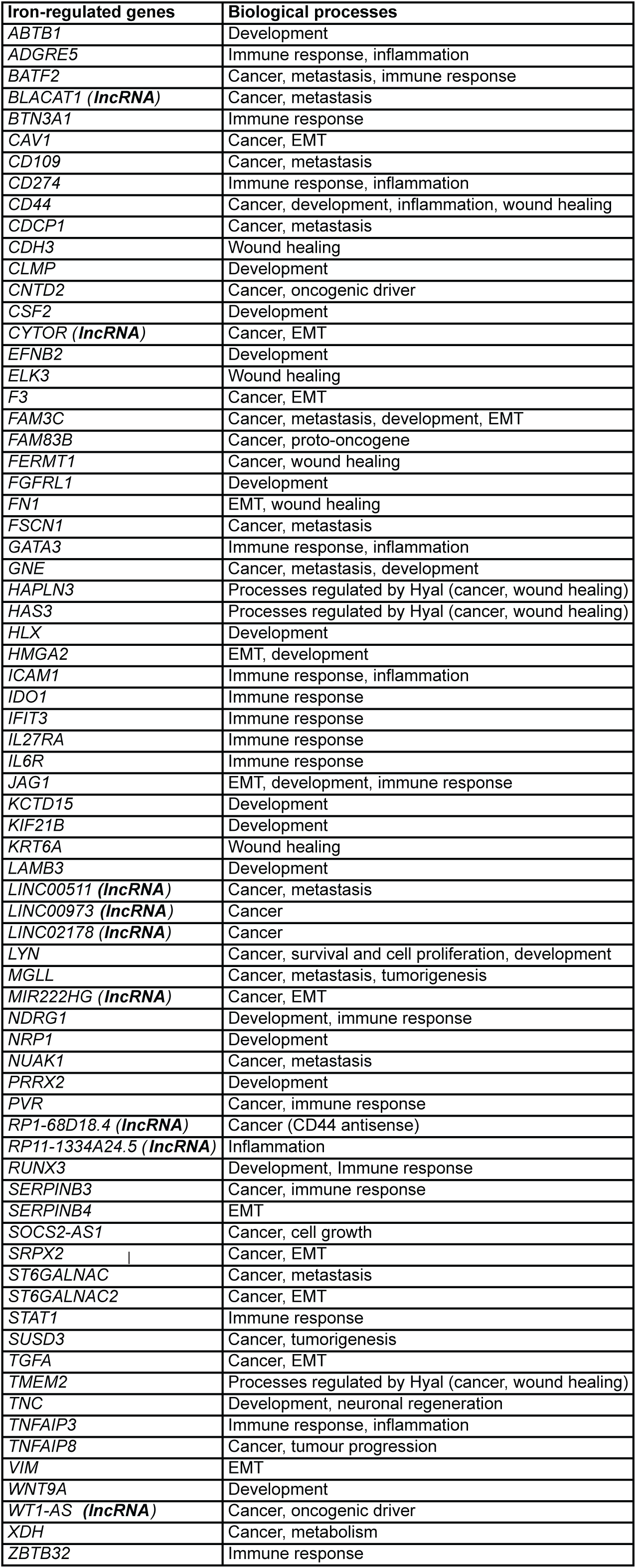
Subset of iron-regulated genes in the mesenchymal state of cells. Related to Figure 4. List of selected genes identified by ChIP-seq where reduction of H3K9me2 reads count correlates with the upregulation of RNA transcripts. CD44 has previously been linked to biological processes implicating iron-regulated genes. These processes include cancer progression, EMT, development, immune responses, inflammation and wound healing.

**Figure S5.**
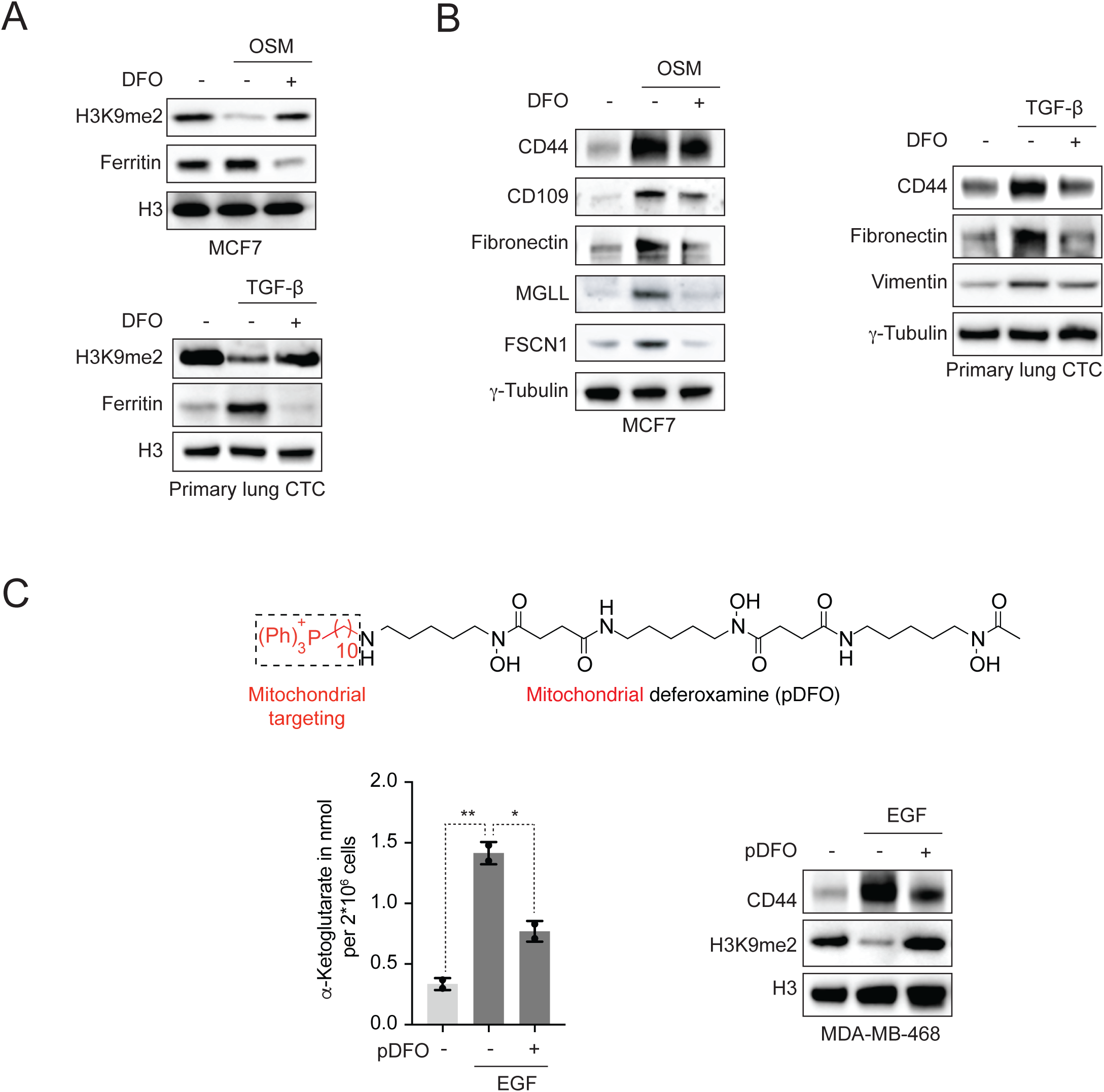
Pharmacological targeting of iron-regulated processes. Related to Figure 5. (**A**)Western blot analysis of H3K9me2 levels in cells cotreated with EGF and DFO. Data showing that targeting nuclear iron antagonizes the effect of EGF, preventing the reduction of H3K9me2. (**B**) Western blot analysis of proteins whose genes are regulated by H3K9me2 in cells cotreated with EGF and DFO. Data showing that targeting nuclear iron antagonizes the effect of EGF, preventing the upregulation of iron-regulated genes. (**C**)Molecular structure of pDFO (top), titration of αKG (bottom left, *n* = 2 technical replicates) and western blot analysis of H3K9me2 levels (bottom right) in cells cotreated with EGF and pDFO. Data showing that targeting mitochondrial iron antagonizes the effect of EGF, blocking the production of αKG, the reduction of H3K9me2 and the upregulation of CD44.

## METHODS

### LEAD CONTACT AND MATERIALS AVAILABILITY

Further information and requests for resources and reagents should be directed to and will be fulfilled by the Lead Contact, Raphaël Rodriguez.

### EXPERIMENTAL MODEL AND SUBJECT DETAILS

#### Cell culture

MCF7 (ATCC, HTB-22, sex: female) and MDA-MB-468 (ATCC, HTB-132, sex: female) cells were grown in Dulbecco’s Modified Eagle Medium GlutaMAX (DMEM, Thermo Fisher Scientific, #61965059) supplemented with 10% Fetal Bovine Serum (FBS, GIBCO, #10270-106) and PEN-STREP (BioWhittaker/Lonza, #DE17-602E) unless stated otherwise. HMLER cells (sex: female) naturally repressing E-cadherin, obtained from human mammary epithelial cells infected with a retrovirus carrying hTERT, SV40 and the oncogenic allele *HrasV12*, were cultured in DMEM/F12 (GIBCO, #31331-028) supplemented with 10% FBS, 10 μg/mL insulin (Sigma-Aldrich, #I0516), 0.5 μg/mL hydrocortisone (Sigma-Aldrich, #H0888) and 0.5 μg/mL puromycin (Life Technologies, #A11138-02), unless stated otherwise. HMLER CD44^high^ cells were also supplemented with 10 ng/mL EGF, whereas HMLER CD44^low^ cells were grown without EGF. HMLER CD44^low/high^ cells stained with CD24-APC and CD44-PE antibodies were sorted by FACS using an Aria IIu (BD Biosciences) to obtain isolated CD44^high^/CD24^low^ and CD44^low^/CD44^high^ cell populations. Primary breast cancer cells (Celther Polska, #RT-04002-CLTH, Lot#074325, sex: female) were grown in BCFM medium (Celther Polska, #MED 02002-CLTH) and kept in culture until the third passage. Primary lung circulating tumor cells (Celprogen, #36107-34CTC, Lot#219411, sex: female) and primary colon circulating tumor cells (Celprogen, #36112-39CTC, Lot#20188, sex: female) were grown using stem cell complete media (Celprogen, #M36102-29PS) until the third passage. Circulating cancer cells were grown in stem cell ECM T75-flasks (Celprogen, # E36102-29-T75) and ECM 6-well plates (Celprogen, #E36102-29-6Well). Primary human T-cells were cultured in RPMI 1640 medium (Gibco, #61870-010) supplemented with 10% FBS. All cells were incubated at 37 °C with 5% CO_2_.

#### Genome editing

CRISPR (Ran et al., 2013) knock out cell lines were generated using the CRISPR plasmid *CD44* (Santa Cruz Biotechnologies, #sc-400209, including the guide RNAs 5’-AATATAACCTGCCGCTTTGC-3’, 5’-TCGCTACAGCATCTCTCGGA-3’ and 5’-CCAGCTATTGTTAACCGTGA-3’) or CRISPR plasmid *TRFC* (Santa Cruz Biotechnologies, #sc-400310, including the guide RNAs 5’-AATTCATATGTCCCTCGTG-3’, 5’-TCTATAGTGATTGTCAGAGC-3’ and 5’-ATCACTATAGATCCATTCAC-3’) and cells were transfected using the TransIT-BrCa Transfection Reagent (Euromedex, #MIR5504) according to the manufacturer’s protocol. Cells were then grown for 3 d and live cells were labeled with CD44-Alexa-Fluor-647 or TfR1-PE antibodies in 2% FBS/1× PBS with 1 mM ETDA and subsequently CD44-negative or TfR1-negative cells were sorted by FACS (Aria IIu, BD Biosciences), seeding one cell per well into 96-well plates. Clones were grown for 14 to 21 d and then analyzed by flow cytometry and western blotting and compared to the parental HMLER CD44^high^ cell line. To validate genomic deletions, genomic DNA was extracted using DNeasy Blood & Tissue Kit (Qiagen, #69504) and amplified by PCR using Phusion Hot Start II DNA Polymerase (Thermo Fisher Scientific, #F549L). Samples were loaded on a 2% agarose gel with a 100 bp DNA ladder (Thermo Fisher Scientific, #15628019). Primers were designed using Primer3-based OligoPerfect (Thermo Fisher Scientific) from the Ensembl genome browser (*CD44*: ENSG00000026508, *TFRC*: ENSG00000072274). Primer sequences (Eurofins Genomics) used were: FWD_*CD44*: 5’-GCCCGGCCTTATTTGACTTT-3’, REV_*CD44*: 5’-GCAGGTCTCAAATCCGATGC-3’; FWD1_*TFRC*: 5’-AAGGCATTTTCTCCTCCAGG-3’, REV1_*TFRC*: 5’-ATGTTGATCACGCCAGACTT-3’; FWD2_*TFRC*: 5’-GTTCCTTGTGTATGTTTGGCTAT-3’, REV2_*TFRC*: 5’-AGGGTAAACTGGTCCATGCT-3’.

## METHODS DETAILS

### Antibody conditions

(WB = Western blot, FC = Flow cytometry, FM = Fluorescence microscopy). CD109 (Santa Cruz Biotechnology, #sc-271085, WB: 1:500), CD24-APC (Sony Biotechnology Inc., #2155590, FC: 1 μL/10^6^ cells), CD44 (Abcam, #ab189524, WB: 1:30000, FM: 1:200), CD44 (Abcam, #ab25340, FC, blocking: 10 μg/mL, 1 h), CD44 Alexa-Fluor-647 (Novus Biologicals, #NB500-481AF647, FC: 1 μL/10^6^ cells), CD44 (R&D Systems, #FAB4948P, FC: 10 μL/10^6^ cells), CDCP1 (Cell Signaling, #4115, WB: 1:1000), *Drosophila* spike-in antibody (Active Motif, #61686, ChIP-seq: 1 µg), E-cadherin (Cell Signaling, #20023195, WB: 1:1000, IF: 1:200), Ferritin (Abcam, #ab75973, WB: 1:1000), Fibronectin (Sigma-Aldrich, #F0791, WB: 1:1000), FSCN1 (Abcam, #ab126772, WB: 1:1000), H3 (Cell Signaling, #9715S, WB: 1:10^6^), H3K9me2 (Cell Signaling, #4658S, WB: 1:1000, ChIP-seq: 6 μL), MGLL (Abcam, #ab24701, WB: 1:1000), PHF8 (Active Motif, #39711, WB: 1:1000), Transferrin receptor 1 (TfR1, Life Technologies, #13-6800, WB: 1:1000), Transferrin receptor 1 (TfR1, Merck Millipore, #GR09L-100UG, FC, blocking: 10 μg/mL, 1 h), TfR1-PE (R&D Systems, #FAB2474P, FC: 5 μL/10^6^ cells), β-Tubulin (Sigma-Aldrich, #T4026-100UL, WB: 1:2000), γ-Tubulin (Sigma-Aldrich, #T5326, WB: 1:2000), Vimentin (Cell Signaling, #3932, WB: 1:500), Zeb1 (Santa Cruz Biotechnology, #sc-81428, WB: 1:500). *Secondary antibodies for WB:* HRP anti-Mouse (Bethyl Laboratories, #A90-116P, 1:10000) and HRP anti-Rabbit (Bethyl Laboratories, #A120-108P, 1:10000). *Secondary antibodies for fluorescence microscopy:* Alexa-Fluor-488 conjugate (Life Technologies, #A-11017 Mouse, #A-11008 Rabbit, 1:1000), Alexa-Fluor-594 conjugate (Life Technologies, #A-11032 Mouse, #A-11072 Rabbit, 1:1000), Alexa-Fluor-647 conjugate (Life Technologies, #A-20990 Mouse, #A-20991 Rabbit, 1:500).

### Reagent conditions

Clickable deferoxamine (cDFO, in-house, 100 μM, 10 min), Deferoxamine (DFO, Sigma-Aldrich, #D9533, 10 μM, 72 h), human epidermal growth factor (Lo et al., 2007) (EGF, Miltenyi Biotech, #130-093-750, 100 ng/mL, 72 h), Fe(III) nitrate (Fe(NO_3_)_3_·9H_2_O, Alfa Aesar, #10715-06), Hyaluronate tetrasaccharide (low-molecular-mass Hyal, TCI Chemicals, #H1284), Hyal 8–15 kDa (Carbosynth, #FH63426, 1 mg/mL), Hyal 40–50 kDa (Carbosynth, #FH01773, 1 mg/mL), Hyal 800 kDa (Carbosynth, #FH45321, 1 mg/mL), Hyaluronate Fluorescein (Hyal-FITC 800 kDa, Carbosynth, #FH45321, 0.1 mg/mL, ∼125 μM), Hyaluronidase (Type IV-S, Sigma-Aldrich, #H3884, 1 mg/mL, 1 h), Ironomycin (in-house, 1 μM, 72 h), Metformin (Alfa Aesar, #J63361, 10 mM, 72 h), Mitochondrial deferoxamine (pDFO, in-house, 500 nM, 72 h), Recombinant oncostatin M (West et al., 2014) (OSM, R&D systems, #295-OM-050, 100 ng/mL, 72 h), Phalloidin-DyLight-488 (Phalloidin-488, Cell Signaling, #12935S, 1:40), RhoNox-M (in-house, 1 μM, 1 h), Transferrin-Alexa-Fluor-647 (Tf-647, Thermo Fisher Scientific, #T23366, 0.01 mg/mL ∼125 μM), Transforming growth factor beta 1 (TGF-β, Miltenyi Biotec, #130-095-066).

### Chemical synthesis and NMR spectroscopy

Starting materials were purchased from Sigma-Aldrich or TCI and used without further purification. Solvents were dried under standard conditions. Reactions were monitored by thin layer chromatography using aluminium-baked plates from Merck (60 F_254_). TLC plates were visualized by UV or treatment with a ninhydrin solution and heating. Reaction products were purified using a preparative HPLC Quaternary Gradient 2545 equipped with a Photodiode Array detector (Waters) fitted with a reverse phase column (XBridge Prep C18 5μm OBD 30×150 mm). NMR spectroscopy was performed on Bruker 300, 400 or 500 MHz apparatus. Spectra were run in CD_3_OD, D_2_O or CDCl_3_, at 298 K or 310 K. ^1^H chemical shifts δ are expressed in ppm using the residual non-deuterated solvent as internal standard and the coupling constants *J* are specified in Hz. The following abbreviations are used: s, singlet; d, doublet; dd, doublet of doublets; dt, doublet of triplets; t, triplet; quint., quintet; m, multiplet. ^13^C chemical shifts δ are expressed in ppm using the residual non-deuterated solvent as internal standard. High-resolution mass spectra (HRMS) were recorded on a Bruker maXis (Q-TOF) mass spectrometer equipped with an ESI ionization source and a TOF detector or a Thermo Fisher Scientific Q Exactive Plus equipped with a Robotic TriVersa NanoMate Advion. Low-resolution mass spectra were recorded on a Waters Acquity H equipped with an SQ Detector 2 (UPLC-MS).

*RhoNox-M* was synthesized in 3 steps according to a previously published procedure (Niwa et al., 2014). ^1^H NMR (300 MHz, CDCl_3_) δ 7.93 (1H, d, *J* = 2.0 Hz), 7.45 (1H, dd, *J* = 8.5 Hz, 2.0 Hz), 7.40–7.30 (3H, m), 7.05 (1H, d, *J* = 8.5 Hz), 6.90 (1H, d, 7.0 Hz), 6.80 (1H, d, *J* = 8.0 Hz), 6.50–6.44 (2H, m), 5.28–5.35 (2H, m), 3.62 (6H, m), 2.97 (6H, s). MS (ESI) m/z: calcd. for C_24_H_25_N_2_O_3_ [M+H]^+^ 389.19, found: 389.35.

#### Clickable deferoxamine (cDFO)

Propargyl bromide (8.50 μL, 0.076 mmol, 80% in toluene) was added to a suspension of DFO mesylate (50 mg, 0.076 mmol) and NaHCO_3_ (0.0192 mg, 0.2284 mmol) in 1 mL of anhydrous MeOH at 0 °C and the reaction mixture was stirred overnight at room temperature. After filtration and concentration under reduced pressure, the crude residue was purified using a preparative HPLC equipped with a C18 reverse phase column (gradient acetonitrile/water/trifluoroacetic acid 5:95:0.1 to acetonitrile/water/trifluoroacetic acid 100:0:0.1) to afford 9.2 mg (17%) of cDFO as a white solid (TFA salt). ^1^H NMR (500 MHz, CD_3_OD) δ 3.94 (2H, d, *J* = 2.5 Hz), 3.71–3.53 (6H, m), 3.25 (1H, t, *J* = 2.5 Hz), 3.17 (4H, t, *J* = 7.0 Hz), 3.08 (2H, t, *J* = 7.5 Hz), 2.76 (4H, t, *J* = 7.0 Hz), 2.46 (4H, dt, *J* = 10.0, 7.0 Hz), 2.09 (3H, s), 1.74–1.61 (8H, m), 1.53 (4H, quint., *J* = 7.0 Hz), 1.46–1.27 (6H, m). ^13^C NMR (125 MHz, CD_3_OD) δ 174.9 (2C), 174.7, 174.5, 173.5, 162.8 (q, *J* = 35.7 Hz), 118.1 (q, *J* = 292.9 Hz), 79.2, 74.6, 48.7 (2C), 48.3, 48.0, 40.3 (2C), 37.3, 31.4, 31.2, 30.0 (2C), 28.9, 28.7, 27.3 (2C), 26.9, 26.4, 24.9 (2C), 24.3, 20.2. HRMS (ESI-TOF) m/z: calcd. For C_28_H_51_N_6_O_8_ [M+H]^+^ 599.3768, found: 599.3760.

#### Triphenylphosphonium deferoxamine (pDFO)

DFO mesylate (100 mg, 0.152 mmol) and NaHCO_3_ (38.3 mg, 0.456 mmol) were added to a solution of (10-bromodecyl)triphenylphosphonium bromide (85.5 mg, 0.152 mmol) in 2 mL of MeOH. The reaction mixture was stirred at 50 °C for 4 h. Then, the solvent was removed under reduced pressure and the residue was purified using a preparative HPLC equipped with a C18 reverse phase column (gradient acetonitrile/water/formic acid 5:95:0.1 to acetonitrile/water/formic acid 100:0:0.1) to afford 12.0 mg (8%) of pDFO as a colorless waxy solid.^1^H NMR (500 MHz, CD_3_OD) δ 8.54 (s, 2H), 7.94–7.86 (m, 3H), 7.84–7.70 (m, 12H), 3.96–3.83 (m, 2H), 3.72–3.53 (m, 6H), 3.47–3.34 (m, 2H), 3.23–3.10 (m, 4H), 3.04–2.85 (m, 2H), 2.85–2.71 (m, 4H), 2.53–2.40 (m, 4H), 2.15–2.04 (m, 3H), 1.81–1.48 (m, 18H), 1.46–1.15 (m, 16H). HRMS (ESI) m/z: calcd. for C_53_H_82_N_6_O_8_P [M]^+^ 961.5926, found: 961.5921.

#### NMR analysis of Hyal/Fe(III) organometallic complexes

In an NMR tube, 1 mol equiv. of a solution of Fe(NO_3_)_3_·9H_2_O in D_2_O/^*t*^BuOH (0.5%) was added to a 2 mM solution of low-molecular-mass Hyal in D_2_O/^*t*^BuOH (0.5%). Then, a drop of TFA was added. In a separate NMR tube, a drop of TFA was added to a 2 mM solution of low-molecular-mass Hyal in D_2_O/^*t*^BuOH (0.5%). NMR spectra were recorded at 310 K on a 500 MHz apparatus.

### Western blotting (WB)

Cells were treated as indicated and then washed with 1× PBS. Proteins were solubilized in 2× Laemmli buffer containing benzonase (VWR, #70664-3, 1:100), extracts were incubated at 37 °C for 1 h, and quantified using a NanoDrop 2000 spectrophotometer (Thermo Fisher Scientific). Protein lysates were resolved by SDS-PAGE electrophoresis (Invitrogen sure-lock system and Nu-PAGE 4–12% Bis-Tris precast gels) and transferred onto nitrocellulose (Amersham Protran 0.45 μm) membranes using a Trans-Blot SD semi-dry electrophoretic transfer cell (Bio-rad). Membranes were blocked with 5% non-fat skimmed milk powder in 0.1% Tween-20/1× PBS for 1 h. Blots were then probed with the relevant primary antibodies in 5% BSA, 0.1% Tween-20/1× PBS at 4 °C overnight with gentle motion. Membranes were washed with 0.1% Tween-20/1× PBS three times and incubated with horseradish peroxidase conjugated secondary antibodies (Jackson Laboratories) in 5% non-fat skimmed milk powder, 0.1% Tween-20/1× PBS for 1 h at room temperature and washed three times with 0.1% Tween-20/1× PBS. Antigens were detected using the SuperSignal West Pico PLUS or West Femto enhanced chemiluminescent detection kits (Thermo Fisher Scientific, #34580 and #34096). Signals were recorded using a Fusion Solo S Imaging System (Vilber) and quantified as indicated using ImageJ.

#### Analysis of ferritin levels upon Hyal treatment

Cells were plated 24 h prior to the experiment. Cells at medium confluence were incubated in FBS-free medium for 2 h. Subsequently, cells were incubated in normal cell culture medium containing Hyal (40–50 kDa) for 24 h. Then, cells were washed with 1× PBS and analyzed by western blotting.

### Fluorescence microscopy (FM)

#### Immunofluorescence

Cells were plated on cover slips 24 h prior to the experiment. Subsequently, cells were washed three times with 1× PBS and fixed with 2% paraformaldehyde in 1× PBS for 12 min, and then washed three times with 1× PBS. After fixation, cells were permeabilized with 0.1% Triton X-100 in 1× PBS for 5 min and washed three times with 1× PBS. Then, cells were blocked in 2% BSA, 0.2% Tween-20/1× PBS (blocking buffer) for 20 min at room temperature. Cells were incubated with the relevant antibody in blocking buffer for 1 h at room temperature and washed three times with blocking buffer. Then, cells were incubated with secondary antibodies for 1 h. Finally, cover slips were washed three times with 1× PBS and mounted using VECTASHIELD (Vector Laboratories) containing DAPI.

#### Other fluorescence imaging experiments

Cells were plated on cover slips 24 h prior to the experiment. Then, cells were grown for 1 h in the appropriate medium supplemented with 0.5% FBS. Live cells were then incubated for 1 h in medium containing RhoNox-M, Hyal-FITC and Tf-647. To this end, cover slips were inverted onto the solutions containing the appropriate reagents and probes in a dish coated with PARAFILM (#PM-996) and incubated at 37 °C with 5% CO_2_. Cover slips were subsequently washed three times with 1× PBS, and mounted using VECTASHIELD containing DAPI. Fluorescence images were acquired using a Deltavision real-time microscope (Applied Precision). 40×/1.4NA, 60×/1.4NA and 100×/1.4NA objectives were used for 2D and 3D acquisitions. Data were deconvoluted with SoftWorx (Ratio conservative −15 iterations, Applied Precision) and processed with ImageJ. All images were acquired as z-stacks.

### Flow cytometry (FC)

Fluorescent probes were used as described in the Reagents section. For surface staining with antibodies, cells were suspended in ice-cold 1× PBS containing 2% FBS and 1 mM EDTA (incubation buffer) and were incubated for 20 min at 4 °C with the relevant antibody. Cells were then washed twice with ice-cold 1× PBS and suspended in incubation buffer prior to being analyzed by flow cytometry. For each condition, at least 10000 cells were counted. Data were recorded on a BD Accuri C6 (BD Biosciences) and processed using Cell Quest (BD Biosciences) and FlowJo (FLOWJO, LLC).

### Inductively coupled plasma mass spectrometry (ICP-MS)

Glass vials equipped with Teflon septa were cleaned with nitric acid 65% (VWR, Suprapur, 1.00441.0250), washed with ultrapure water (Sigma-Aldrich, #1012620500) and dried. Cells were plated 24 h prior to the experiments. In all experiments, cells were incubated for 2 h with FBS-free medium prior to treatment. Cells were harvested using trypsinization (TrypLE Express Enzyme, Life Technologies, 12605010) followed by two washes with 1× PBS. Cells were then counted using an automated cell counter (Entek) and transferred in 100 µL 1× PBS to clean vials, and samples were lyophilized using a freeze dryer (CHRIST, #22080). Samples were subsequently mixed with nitric acid 65% overnight followed by heating at 80 °C for 2 h. Samples were diluted with ultrapure water (Sigma-Aldrich, 1012620500) to a final concentration of 0.475 N nitric acid and transferred to metal-free centrifuge vials (VWR, 89049-172) for subsequent ICP-MS analysis. ^56^Fe concentrations were measured using an Agilent 7900 ICP-QMS in low-resolution mode. Sample introduction was achieved with a micro-nebulizer (MicroMist, 0.2 mL/min) through a Scott spray chamber. Isotopes were measured using a collision-reaction interface with helium gas (5 mL/min) to remove polyatomic interferences. Scandium and indium internal standards were injected after inline mixing with the samples to control the absence of signal drift and matrix effects. A mix of certified standards was measured at concentrations spanning those of the samples to convert count measurements to concentrations in the solution. Uncertainties on sample concentrations were calculated using algebraic propagation of ICP-MS blank and sample counts uncertainties. Values were normalized by dry weight and cell number.

### Transfection

HMLER CD44^high^ and *CD44* ko2 cells were plated 24 h prior to the experiment. *CD44* ko2 cells were transfected with a *CD44* cDNA plasmid (Interchim, #HG12211-UT) using TransIT-BrCa Transfection reagent according to the manufacturer’s protocol, and control cells were treated using TransIT-BrCa Transfection reagent without DNA. The medium was changed 24 h post transfection and cells incubated for 48 h. Subsequently, cells were incubated for 1 h in medium without FBS. Imaging of RhoNox-M and Hyal-FITC was performed and analyzed as described in the Fluorescence microscopy section.

### RNA interference

Cells were plated 24 h prior to the experiments. Cells were transfected with the specified siRNA using jetPRIME (Polyplus, #114-15) according to the manufacturer’s protocol in DMEM supplemented with PEN-STREP and 10% FBS. Cells were then washed with 1× PBS and harvested using 2× Laemmli buffer. Suitable small interfering RNAs were designed by Dharmacon for specific down-regulation of CD44 (ON-TARGETplus *CD44* siRNA SMARTpool, #L-009999-09-0020, target sequences: 5′-GAAUAUAACCUGCCGCUUU-3′, 5′-CAAGUGGACUCAACGGAGA-3′, 5′-CGAAGAAGGUGUGUGGGCAGA-3′ and 5′-GAUCAACAGUGGCAAUGGA-3′), TfR1 (ON-TARGETplus *TRFC* siRNA, SMARTpool, #L-003941-00-0005, target sequences: 5′-GAAUGGAUCUAUAGUGAUU-3′, 5′-GAUAAGAACGGUAGACUUG-3′, 5′-GUAAACUGGUCCAUGCUAA-3′, 5′-CUGAAUGGCUAGAGGGAUA-3′) and PHF8 (ON-TARGETplus *PHF8* siRNA SMARTpool, L-004291-01-0005, target sequences: 5′-CUCAUGAGUGUGCGAGAUA-3′, 5′-UCAAGAAGGCAGAGCGAAA-3′, 5′-GGUGAUGGAAGACGAAUUU-3′ and 5′-UGGGAGUGUUAGUAAUCAA-3′). Control siRNA (Qiagen, #1027310) was used for comparison.

### RT-qPCR

Total RNA was extracted using the miRNeasy mini kit (QIAGEN) according to the manufacturer’s protocol. RNA was quantified using a NanoDrop 2000 spectrophotometer (Thermo Fisher Scientific). 2 µg of total RNA was used to synthesize cDNA using oligo(dT) primers and the SuperScript II Reverse Transcriptase (Invitrogen). cDNA was quantified by qPCR using SYBR Green 1 Master mix (Roche) on a LightCycler 480 (Roche) real-time PCR system. The data were normalized against *GAPDH* or *RPL11* housekeeping genes. Primer sequences were generated using Primer-BLAST (https://www.ncbi.nlm.nih.gov/tools/primer-blast/). Primer sequences used were: FWD_*CD44*: 5’-CCCTGCTACCAATAGGAATGATG-3’, REV_*CD44*: 5’-CTGCTTTCCTTCGTGTGTGG-3’; FWD_*TFRC*: 5’-ATCGGTTGGTGCCACTGAAT-3’, REV_*TFRC*: 5’-TTGCTGGTACCAAGAACCGC-3’; FWD_*FTH1*: 5’-CCAGAACTACCACCAGGACT-3’, REV_*FTH1*: 5’-GGTCAAAGTAGTAAGACATGGACAG-3’; FWD_*GAPDH*: 5’-TCACCATCTTCCAGGAGCGA-3’, REV_*GAPDH*: 5’-GATGACCCTTTTGGCTCCCC-3’; FWD_*RPL11*: 5’-AGCAGCCAAGGTGTTGGAG-3’, REV_*RPL11*: 5’-TACTCCCGCACCTTTAGACC-3’.

### Proteomics

Cells were grown in 148 cm^2^ round dishes and treated as described in the manuscript. Whole cell extracts were collected and then centrifuged at 500 × g for 5 min at 4 °C, washed twice in ice-cold 1× PBS and lysed using lysis buffer (8 M urea, 50 mM NH_4_HCO_3_ and cOmplete (Roche, #000000011697498001)). The global proteome was quantitatively analyzed with a Q Exactive HF-X quadrupole-Orbitrap mass spectrometer using a label-free approach. About 30 µg of total protein cell lysate was diluted with 90 µL of 25 mM NH_4_HCO_3_. Protein was reduced by incubation with 5 mM dithiothreitol (DTT) at 57 °C for 1 h and then alkylated with 9 mM iodoacetamide for 30 min at room temperature in the dark. Trypsin/LysC (Promega) was added at 1:100 (wt:wt) enzyme:substrate. Digestion was performed overnight at 37 °C. Samples were then loaded onto a homemade C18 StageTips for desalting. Peptides were eluted from beads by incubation with 40:60 MeCN/H_2_O with 0.1% formic acid. Peptides were dried in a Speedvac and reconstituted in 15 µL 2:98 MeCN/H_2_O with 0.3% TFA prior to liquid chromatography-tandem mass spectrometry (LC-MS/MS) analysis. Samples (4 µL) were chromatographically separated using an RSLCnano system (Ultimate 3000, Thermo Scientific) coupled online to a Q Exactive HF-X with a Nanospray Flex ion source (Thermo Scientific). Peptides were first loaded onto a C18-trapped column (75 μm inner diameter × 2 cm; nanoViper Acclaim PepMap 100, Thermo Scientific), with buffer A (2:98 MeCN/H_2_O with 0.1% formic acid) at a flow rate of 2.5 µL/min over 4 min and then switched for separation to a C18 column (75 μm inner diameter × 50 cm; nanoViper C18, 2 μm, 100 Å, Acclaim PepMap RSLC, Thermo Scientific) regulated to a temperature of 50 °C with a linear gradient of 2 to 35% buffer B (100% MeCN and 0.1% formic acid) at a flow rate of 300 nL/min over 211 min. MS full scans were performed in the ultrahigh-field Orbitrap mass analyzer in ranges *m*/*z* 375–1500 with a resolution of 120000 at *m*/*z* 200, the maximum injection time (MIT) was 50 ms and the automatic gain control (AGC) was set to 3 × 10^6^. The top 20 intense ions were subjected to Orbitrap for further fragmentation via high energy collision dissociation (HCD) activation and a resolution of 15000 with the intensity threshold kept at 3.2 × 10^5^. We selected ions with charge state from 2+ to 6+ for screening. Normalized collision energy (NCE) was set at 27. For each scan, the AGC was set at 1 × 10^5^, the MIT was set at 25 ms and a dynamic exclusion of 40 s. The identity of proteins was established from UniProt human canonical database (downloaded on 22/08/2017) using Sequest HF through proteome discoverer (version 2.0). Enzyme specificity was set to trypsin and a maximum of two missed cleavage sites were allowed. Oxidized methionine, N-terminal acetylation, and carbamidomethyl cysteine were set as variable modifications. Maximum allowed mass deviation was set to 10 ppm for monoisotopic precursor ions and 0.02 Da for MS/MS peaks. The resulting files were further processed using myProMS v3.6 (Poullet et al., 2007). FDR calculation used Percolator and was set to 1% at the peptide level for the whole study. The label-free quantification was performed by peptide Extracted Ion Chromatograms (XICs) computed with MassChroQ version 2.2.2 (Valot et al., 2011). For protein quantification, XICs from proteotypic peptides shared between compared conditions (TopN matching) with up to two missed cleavages and carbamidomethyl modifications were used. Median and scale normalization was applied on the total signal to correct the XICs for each biological replicate. To estimate the significance of the change in protein abundance, a linear model (adjusted on peptides and biological replicates) was performed and *P*-values were adjusted with a Benjamini–Hochberg FDR procedure with a control threshold set to 0.05. Fold change-based GO enrichment analysis was performed as in (Kowal et al., 2016).

### Metabolomics

Metabolomics experiments were performed and analyzed as previously described (Pietrocola et al., 2018; Pietrocola et al., 2017). Briefly, for sample preparation, cells were grown in 6-well plates and treated as indicated. Plates were placed on ice, the medium was carefully removed and cells were quickly washed three times with 1 mL ice-cold 1× PBS and then lysed with 1 mL cold lysis buffer (methanol/water, 9:1, v/v, –20 °C). Next, cells were mixed by vortexing for 1 min and centrifuged at 15000 × g for 10 min at 4 °C. For GC-MS analysis of metabolites, 750 µL of the supernatant was transferred to centrifugation microtubes and stored at –80 °C. Supernatants were completely evaporated at 40 °C. Next, 500 μL methanol was added to dried extracts, 270 µL was transferred to injection vials and evaporated at 40 °C, 50 µL of methoxyamine (20 mg/mL in pyridine) was added to dried extracts and samples were stored at room temperature in the dark for 16 h. Then, 80 µL of MSTFA was added and final derivatization occurred at 40 °C for 30 min. Widely-targeted analysis of intracellular metabolites was performed using gas chromatography (GC) coupled to a triple quadrupole (QQQ) mass spectrometer. The GC-MS/MS method was performed on a 7890B gas chromatography (Agilent Technologies) coupled to a triple quadrupole 7000C (Agilent Technologies) equipped with a high sensitivity electronic impact source (EI) operating in positive mode. The front inlet temperature was 250 °C and the injection was performed in splitless mode. The transfer line and the ion-source temperature were 250 °C and 230 °C, respectively. The septum purge flow was fixed at 3 mL/min, the purge flow to split vent operated at 80 mL/min for 1 min and gas saver mode was set to 15 mL/min after 5 min. The helium gas flowed through the column (J&WScientificHP-5MS, 30 m × 0.25 mm, i.d. 0.25 mm, d.f., Agilent Technologies) at 1 mL/min. The column temperature was held at 60 °C for 1 min, then raised to 210 °C (10 °C/min), followed by a step at 230 °C (5 °C/min), then reached 325 °C (15 °C/min) and was maintained at this temperature for 5 min. Nitrogen was used as collision gas. The MRM for biological samples was used as scan mode. Peak detection and integration of each analyte were performed using the Agilent Mass Hunter quantitative software (B.07.01).

### Titration of α-ketoglutarate

Levels of αKG were quantified using an αKG assay kit (Abcam, #ab83431) according to the manufacturer’s protocol. Cells were grown in 60.1 cm^2^ round dishes and treated as indicated. Cells were then washed with ice-cold 1× PBS, harvested by scraping and resuspended in ice-cold αKG buffer (kit component). Cells were centrifuged at 25000 × g for 5 min at 4 °C and the supernatant was transferred to clean tubes. Ice-cold perchloric acid was added to a final concentration of 1 M and the solution was incubated on ice for 5 min. Cells were centrifuged at 13000 × g for 2 min and the supernatant was transferred to a clean tube. Then, 1/3 vol. of a 2 M solution of KOH was added and the pH adjusted to 7.4 using a 0.1 M aqueous solution of KOH. Samples were centrifuged at 13000 × g for 15 min and the supernatants collected. αKG levels were measured according to the manufacturer’s protocol, using a standard curve and a control to subtract pyruvate background levels. Fluorescence intensities (ex. 535 nm; em. 590 nm) were recorded using a Perkin Elmer Wallac 1420 Victor2 Microplate Reader, and data were normalized against cell numbers. Values were derived from the standard curve.

### Subcellular fractionation

Nuclear and cytoplasmic fractions were isolated using the subcellular fractionation kit for cultured cells according to the manufacturer’s protocol (Thermo Fisher Scientific, #78840). Cells were cultured in round dishes, harvested using trypsin-EDTA and washed in ice-cold 1× PBS. Combining the cytosolic and membrane fractions yielded the cytoplasmic fraction, while combining the nuclear soluble and chromatin-associated fractions generated the nuclear fraction.

### Histone extraction

Cells were grown in 148 cm^2^ round dishes and treated as indicated. Whole cell extracts were collected by scraping. After centrifugation at 500 × g, 4 °C for 5 min, cells were washed with ice-cold 1× PBS. Subsequently, cells were lysed in hypotonic lysis buffer (10 mM HEPES, 10 mM KCl, 1.5 mM MgCl_2_, 0.5 mM DTT, cOmplete) for 45 min at 4 °C on a rotary wheel. After centrifugation at 20000 × g, 4 °C for 20 min, supernatants were discarded and nuclei were resuspended in 0.2 M H_2_SO_4_ overnight at 4 °C on a rotary wheel. Subsequently, lysates were centrifuged at 20000 × g, 4 °C for 20 min and histones were precipitated using 33% trichloroacetic acid for 1 h, at 4 °C. The resulting solutions were then centrifuged at 20000 × g, 4 °C for 20 min to precipitate histones, washed twice with ice-cold acetone and centrifuged at 20000 × g, 4 °C for 10 min. Histones were dried for 20 min at room temperature and solubilized in deionized water. For subsequent mass spectrometry analysis, histones were resolved on an 18% SDS-PAGE, followed by colloidal blue staining.

### Quantitative mass spectrometry by Parallel-Reaction Monitoring (PRM)

Histones were separated by SDS-PAGE and stained with colloidal blue (LabSafe Gel Blue GBiosciences). One gel slice was excised for each purification and in-gel digested using trypsin/LysC (Promega). Peptides extracted from each band were then analyzed by nanoLC-MS/MS (RSLCnano system coupled to a Q Exactive HF-X). Peptides were first trapped onto a C18 column (75 μm inner diameter × 2 cm; nanoViper Acclaim PepMapTM 100, Thermo Scientific) with buffer A at a flow rate of 2.5 µL/min over 4 min. Separation was performed on a 50 cm × 75 µm C18 column (nanoViper C18, 3 μm, 100 Å, Acclaim PepMapTM RSLC, Thermo Scientific) regulated to 50 °C and with a linear gradient from 2% to 35% buffer B at a flow rate of 300 nL/min over 91 min. The mass spectrometer was operated in PRM mode (conditions listed thereafter). The acquisition list was generated from the peptides obtained from mixed samples of 5 biological replicates for each condition based on the data-dependent acquisition results.

| Peptide sequence and modification position | Protein modification | Precursor mass MH+ | Charge | Extracted fragments |
| --- | --- | --- | --- | --- |
| K[Dimethyl (K)]STGGK[Acetyl (K)]APR | H3K9me2 | 486,285265 | 2 | y4, y5, y6, y7, y8, y7++, b6, b7, b8, b6++ |
|  |  | 324,525936 | 3 | y4, y5, y6, y4++, b3, b4, b5, b6 |
| K[Trimethyl (K)]STGGK[Acetyl (K)]APR | H3K9me3 | 493,29309 | 2 | y4, y5, y6, y7, y6++, b8 |
| K[Trimethyl (K)]SAPATGGVK | H3K27me3 | 479,290016 | 2 | y4, y5, y7, y8, y9, y7++, b5, b6, b8, b9, b8++ |
| STELLIR |  | 416,25036 | 2 | y3, y4, y5, y6, y3++, y4++, y6++, b3, b4, b4++ |
| YRPGTVALR |  | 516,801083 | 2 | y3, y4, y5, y6, y7, y7++, y8++, b5, b6, b7, b8, b7++ |
|  |  | 344,869814 | 3 | y3, y4, y5, y6, y3++, y4++, y7++, y8++, b4, b5, b6, b5++ |

The PRM data were analyzed using Skyline (version 4.1, MacCoss Lab Software, Seattle, WA); (https://skyline.ms/project/home/software/Skyline/begin.view), fragment ions for each targeted mass were extracted and peak areas were integrated. The peptide areas were log2 transformed and the mean log2-area was normalized by the mean area of peptide STELLIR and YRPGTVALR using software R (version 3.1.0).

### ChIP-seq

Cells were grown in 148 cm^2^ round dishes and treated as described. Cells were harvested by trypsinization. Medium was added to the cell suspension and cells were centrifuged at 500 × g for 5 min at room temperature. The pelleted cells were resuspended in medium, counted and crosslinked with 1% formaldehyde for 10 min at room temperature. Then, 2.5 M glycine was added to a final concentration of 0.125 M and incubated for 5 min at room temperature followed by centrifugation at 500 × g for 5 min at 4 °C. The pelleted cells were washed twice with ice-cold 1× PBS and collected by centrifugation at 500 × g for 5 min at 4 °C. Pellets were resuspended in lysis buffer A (50 mM Tris-HCl pH 8, 10 mM EDTA, 1% SDS, cOmplete) at room temperature and incubated for 5 min on a rotating wheel. Next, lysates were centrifuged at 300 × g for 10 min at 8 °C to prevent SDS precipitation and supernatants were discarded. Pellets were then sheared in buffer B (25 mM Tris-HCl pH 8, 3 mM EDTA, 0.1% SDS, 1% Triton X-100, 150 mM NaCl, cOmplete) to approximately 200–600 base-pair average size using a Bioruptor Pico (Diagenode). After centrifugation at 20000 × g, 4 °C for 15 min, supernatants containing sheared chromatin were used for immunoprecipitation. 25 µL (10%) of sheared chromatin was used as input DNA to normalize sequencing data. As an additional control for normalization, spike-in chromatin from *Drosophila* and a spike-in antibody were used. Chromatin immunoprecipitation (1 million cells per ChIP) was carried out using sheared chromatin and an antibody against H3K9me2 subsequently complexed to Dynal Protein G magnetic beads (Thermo Fisher). Briefly, 6 µL of H3K9me2 antibody were mixed with 1 µg of spike-in antibody. 22 µL magnetic beads were washed three times in ice-cold buffer C (20 mM Tris-HCl pH 8, 2 mM EDTA, 0.1% SDS, 1% Triton X-100, 150 mM NaCl, cOmplete) and incubated with the mixture of H3K9me2 and spike-in antibody for 4 h, at room temperature on a rotating wheel in buffer C (494 µL). After spinning and removal of supernatants, beads were resuspended in 50 µL buffer C. This suspension was subsequently incubated with 250 µL sheared chromatin previously mixed with 50 ng of spike-in chromatin from *Drosophila* (250 µL chromatin of interest: 2.5 µL spike-in chromatin) at 4 °C on a rotating wheel overnight (16 h). After a spinning, supernatants were discarded and beads were successively washed in buffer C, buffer D (20 mM Tris-HCl pH 8, 2 mM EDTA, 0.1% SDS, 1% Triton X-100, 500 mM NaCl), buffer E (10 mM Tris-HCl pH 8, 0.25 M LiCl, 0.5% NP-40, 0.5% sodium deoxycholate, 1 mM EDTA) and buffer F (10 mM Tris-HCl pH 8, 1 mM EDTA, 50 mM NaCl). Finally, input and immunoprecipitated chromatin samples were resuspended in a solution containing TE buffer/1% SDS, de-crosslinked by heating at 65 °C overnight and subjected to both RNase A (Thermo Fisher Scientific, #12091-039, 1 mg/mL) and Proteinase K (Thermo Fisher Scientific, #E00491, 20 mg/mL) treatments. *Input and immunoprecipitated DNA extraction*: After reverse crosslinking, input and immunoprecipitated chromatin samples were treated with RNase A and proteinase K, and glycogen (Thermo Fisher Scientific, #R0561, 20 mg/mL) was added. Samples were incubated at 37 °C for 2 h. DNA precipitation was carried out using 8 M LiCl (final concentration 0.44 M) and phenol:chloroform:isoamyl alcohol. Samples were vortexed and centrifuged at 20000 × g for 15 min at 4 °C. The upper phase was mixed with chloroform by vortexing. After centrifugation at 20000 × g for 15 min at 4 °C, the upper phase was mixed with –20 °C absolute ethanol by vortexing and stored at –80 °C for 2 h. Next, samples were pelleted at 20000 × g for 20 min at 4 °C. The pellets were washed with ice-cold 70% ethanol and centrifuged at 20000 × g for 15 min at 4 °C. The supernatants were discarded and pellets were dried at room temperature, dissolved in nuclease-free water and quantified using a Qubit fluorimetric assay (Thermo Fisher Scientific) according to the manufacturer’s protocol. *Library preparation and Sequencing*: Illumina compatible libraries were prepared from input and immunoprecipitated DNAs using the Illumina TruSeq ChIP library preparation kit according to the manufacturer’s protocol. Briefly, 4 to 10 ng of DNA was subjected to end-repair, dA-tailing and ligation of TruSeq indexed Illumina adapters. After a final PCR amplification step, the resulting barcoded libraries were equimolarly pooled into 2 groups quantified using a qPCR method (KAPA library quantification kit, Roche) before sequencing on the Illumina HiSeq 2500 instrument. Each pool (11 pM) was loaded on a rapid flow cell and sequenced using a single read mode (SR50). This sequencing configuration was set to reach an average of 25 million reads (50 base pairs long) per sample.

### RNA-seq

MDA-MB-468 cells were grown in 148 cm^2^ round dishes and treated with EGF for 72 h. Whole cell extracts (∼9 × 10^5^ cells) were collected by centrifugation at 500 × g for 5 min at 4 °C and washed twice with ice-cold 1× PBS. Total RNAs were extracted using an RNeasy Mini Kit (Qiagen) according to the manufacturer’s protocol and subjected to DNase I treatment using the RNase-free DNase set (Qiagen). Total RNA was then analyzed using an Agilent RNA 6000 Nano Kit on the Agilent 2100 Bioanalyzer System to determine RNA concentration, RNA Integrity Number (RIN), which was high with RIN = 10 for all samples, and to confirm removal of DNA contamination. After quality control of RNA, the ERCC RNA Spike-In (Ambion) was added to RNA samples (RNA samples:RNA spike-in control diluted at 1/100, 5:1). RNA sequencing libraries were prepared from 1 µg total RNA using the Illumina TruSeq Stranded mRNA Library Preparation Kit, which allows strand specific sequencing to be performed. A first step of polyA selection using magnetic beads was performed to allow sequencing of polyadenylated transcripts. After fragmentation, cDNA synthesis was performed and resulting fragments were used for dA-tailing followed by ligation of TruSeq indexed adapters. PCR amplification was finally achieved to generate the final barcoded cDNA libraries. The 4 libraries were equimolarly pooled and subjected to qPCR quantification (KAPA library quantification kit, Roche). Sequencing was carried out on a NovaSeq 6000 instrument from Illumina based on a 2*100 cycle mode (paired-end reads, 100 bases) using a S1 flow cell in order to obtain around 35 million clusters (70 million raw paired-end reads) per sample.

### ChIP-seq data analysis

Raw sequencing reads were first aligned with the Bowtie 2 software v2.2.9 on the *Drosophila Melanogaster* (dmel-R6,21) reference genome to extract reads coming from spike-in chromatin (Langmead and Salzberg, 2012). Remaining reads were then aligned with the Human hg38 reference genome using Bowtie 2 very-sensitive mode. PCR duplicates were removed from aligned data using Picard tools, and genome-browser track files were generated using the deepTools2 suite (v3.1.0) and the bamCoverage function (Ramirez et al., 2016). In order to validate the overall H3K9me2 profiles, large genomic domains were called with the EPIC software (v0.2.9, https://github.com/biocore-ntnu/epic) and LOCKs were defined as previously proposed (McDonald et al., 2017; McDonald et al., 2011). Counts of aligned reads per human gene were generated using the FeatureCounts software and the gene coordinates extracted from the GENCODE annotation file (v26) (Liao et al., 2014). Genes with less than 10 reads were discarded from the analysis. The raw count table was then normalized using the TMM method from the edgeR R package (v3.22.3) (Robinson et al., 2010), and the limma voom (v3.36.3) functions were applied to detect genes with differential binding between untreated and EGF samples. Finally, genes with an adjusted *P*-value < 0.05 and a log_2_(fold-change) > 0.3 were called significant.

### RNA-seq data analysis

Raw sequencing reads were aligned on the human hg38 reference genome using the STAR mapper (2.5.3a) (Dobin et al., 2013), up to the generation of a raw count table per gene (Gencode annotation v26). Expressed genes (TPM ≥ 1 in at least one sample) have then been selected for downstream analysis. The raw count table was then normalized using the TMM method from the edgeR R package (v3.22.3) (Robinson et al., 2010), and the limma voom (v3.36.3) functions were applied to detect genes with differential expression between untreated and EGF samples. Genes with an adjusted *P*-value < 0.05 and a log_2_(fold-change) > 1 were called significant.

### Correlative analysis of H3K9me2 ChIP-seq and RNA-seq data

Genes that were significantly less enriched in H3K9me2 and overexpressed in terms of RNA levels, or more enriched and repressed, are represented by H3K9me2↓/RNA↑ and H3K9me2↑/RNA↓, respectively. Genes that were both enriched (or depleted) for H3K9me2 and RNA levels were called H3K9me2↑/RNA↑ and H3K9me2↓/RNA↓, respectively. Genes for which the effect of EGF was not significant in at least one of the two types of experiment were called ‘not significant’.

### Chemical labeling of cDFO in cells

Cells were cultured on cover slips and treated with 100 μM cDFO for 10 min. Cells were fixed with formaldehyde (2% in 1× PBS, 12 min) prior to permeabilization (0.1% Triton X-100 in 1× PBS, 5 min) and washed three times with 1% BSA/1× PBS. The click reaction cocktail was prepared from Click-iT EdU Imaging Kits (Thermo Fisher Scientific, #C10337) according to the manufacturer’s protocol as previously described (Mai et al., 2017). Briefly, 868 μL of 1× Click-iT reaction buffer was mixed with 40 μL CuSO_4_ solution, 2 μL Alexa Fluor azide and 90 μL reaction buffer additive (sodium ascorbate) to reach a final volume of 1 mL. Cover slips were incubated with 50 μL of the click reaction cocktail in the dark at room temperature for 30 min, then washed three times with 1× PBS. Cover slips were mounted using VECTASHIELD containing DAPI and were sealed with nail varnish Express manucure (Maybelline, #16P201).

### Cell Viability

Cell viability was evaluated by plating 1000 cells/well in 96-well plates using CellTiter-Blue viability assay according to the manufacturer’s protocol. Cells were treated with different concentrations of ironomycin as indicated for 72 h. CellTiter-Blue reagent (Promega, G8081) was added after 72 h treatment and cells were incubated for 2 h before recording fluorescence intensities (ex. 560 nm; em. 590 nm) using a Perkin Elmer Wallac 1420 Victor2 Microplate Reader.

## QUANTIFICATION AND STATISTICAL ANALYSIS

### Fluorescence image quantifications

Fluorescence intensity quantifications were performed using pixel intensity divided by area. Phalloidin-488 staining was used to define whole cell area where appropriate. Quantifications of iron-containing vesicles were performed by counting *foci* in one fluorescent channel colocalizing with *foci* in another. For example, the number of RhoNox-M-positive vesicles (red *foci*) that contained either Hyal-FITC (green *foci*) or Tf-647 (purple *foci*) was quantified for each cell. To this end, three independent laser channels were used to detect each of the fluorophores. Fluorescence intensities were normalized against untreated control cells for each channel and experiment, and pixel intensity was defined as zero below this control threshold in each channel. Then, vesicles containing iron (RhoNox-M-positive vesicles) were counted for each cell and defined as Hyal-positive or Tf-positive vesicles if the intensity was above the control threshold. Each vesicle that contained iron and Hyal but not Tf were positive in the red and green channels but remained black in the purple channel (zero-pixel intensity). In contrast, vesicles that contained iron and Tf were positive in the red and purple channels but remained black (zero-pixel intensity) in the green channel.

### Western blot quantifications

Western blot signals were quantified using ImageJ. Equal areas were defined for each lane and quantified using the Plot Lanes function in ImageJ. Signals arising from individual antibodies were normalized against the antibody signal of the loading control.

### Statistical Analysis

All results are presented as mean values ± standard deviation (s.d.) unless stated otherwise. PRISM 8 software was used to calculate significance using unpaired *t*-tests. PRISM 8 software or R programming language was used to generate graphical representations of quantifications unless stated otherwise. *P*-values are defined as: * *P* ≤ 0.05, ** *P* ≤ 0.01, *** *P* ≤ 0.001, **** *P* ≤ 0.0001; n.s., not significant. Sample sizes (*n*) are indicated in the figure legends. For PRM: On each peptide, a linear model was used to calculate the mean fold change between the conditions, its 95% confidence interval and the *P*-value of the two-sided associated *t*-tests. *P*-values were adjusted using the Benjamini-Hochberg procedure (Benjamini and Yekutieli, 2001).

## DATA AND CODE AVAILABILITY

### Data availability

The mass spectrometry data have been deposited at the ProteomeXchange Consortium via the PRIDE (Deutsch et al., 2017; Perez-Riverol et al., 2019) partner repository. ChIP-seq and RNA-seq data are available on the National Center for Biotechnology Information website (https://www.ncbi.nlm.nih.gov/geo/query/acc.cgi?acc=GSE121664). Access upon request to the authors. All other data are available from the corresponding author upon request.

### Code availability

Code employed for ChIP-seq and RNA-seq data analyses are available upon request to the authors.

